# Plant diversity darkspots for global collection priorities

**DOI:** 10.1101/2023.09.12.557387

**Authors:** Ian Ondo, Kiran L. Dhanjal-Adams, Samuel Pironon, Daniele Silvestro, Matheus Colli-Silva, Victor Deklerck, Olwen M. Grace, Alexandre K. Monro, Nicky Nicolson, Barnaby Walker, Alexandre Antonelli

**Author notes:** Equal contributions (Ian Ondo, Kiran L. Dhanjal-Adams, Samuel Pironon).

## Abstract

- More than 15% of all vascular plant species may remain scientifically undescribed, and many of the >340,000 described species have no or few geographic records documenting their distribution. Identifying and understanding taxonomic and geographic knowledge shortfalls is key to prioritising future collection and conservation efforts.
- Using extensive data for 343,523 vascular plant species and time-to-event analyses, we conducted multiple tests related to plant taxonomic and geographic data shortfalls, and identified 32 global diversity darkspots (regions predicted to lack most information about their species diversity and distribution). We defined priority regions for future collection according to several socio-economic and environmental scenarios.
- Most plant diversity darkspots are found within biodiversity hotspots, except New Guinea. We identify New Guinea, the Philippines, Borneo, Myanmar, India, Turkey, Colombia, Ecuador, and Peru, as global collection priorities under all environmental and socio-economic conditions considered.
- Our study provides a framework to accelerate plant species documentation for the implementation of conservation actions. As digitisation of the world’s herbaria progresses, collection and conservation priorities may soon be identifiable at finer scales.

## INTRODUCTION

Biodiversity is declining worldwide, posing a direct threat to planetary health (Díaz *et al*., 2019). The recently adopted Kunming-Montreal Global Biodiversity Framework (GBF) aims to facilitate the implementation of biodiversity-positive actions across all levels of society to achieve the vision of “people living in harmony with nature” by 2050. In this context, plant scientists have a key role to play by providing essential data, tools and evidence to decision-makers through improved documentation and understanding of plant diversity. A major scientific milestone in this pursuit is the compilation of names and geographic distributions of nearly 345,000 taxonomically accepted vascular plant species by the World Checklist of Vascular Plants (WCVP; Govaerts *et al*., 2021). The burning questions now are how many species remain to be scientifically described, and where and how to concentrate efforts to safeguard global plant diversity into the future (Pimm & Joppa, 2015).

In an attempt to categorise our knowledge limits, seven biodiversity shortfalls have been described to characterise data gaps (Hortal *et al*., 2015). Two of these shortfalls concern species’ scientific descriptions and geographic distributions, commonly referred to as the Linnaean and Wallacean shortfalls, respectively (Hortal *et al*., 2015). The Linnaean shortfall is assumed to be large, with >60,000 angiosperm species estimated to not have been scientifically named yet (Joppa *et al*., 2011). Moreover, and in contrast to vertebrates (Moura & Jetz, 2021), little is known about the diversity and distribution of undescribed vascular plant species. Although WCVP provides comprehensive distribution data for most known species at a coarse resolution (i.e., “botanical countries”), major biases and gaps have been shown to nurture the Wallacean shortfall at a finer resolution (Meyer *et al*., 2016). This in turn impedes thorough characterisation and understanding of the full geographic range of individual plant species and their diversity.

Assessing the magnitude and spatial variation of both the Linnaean and Wallacean shortfalls is thus key for developing collection strategies. Historical collections have been biased spatially by land accessibility, the density of taxonomists, and both environmental and socio-economic factors (Meyer *et al*., 2016). However, there is no current measure of whether ongoing botanical surveys are being conducted in the places that need them most, or whether they accentuate biases in botanical knowledge (Geldmann *et al*., 2016). It is also important to note that the classification, distribution, and ecology of plants are held extensively in non-western traditional knowledge systems, in which similar shortfalls may not necessarily be quantifiable at large spatial scales (Gardner *et al*., 2022).

Nearly 40% of described vascular plant species are estimated to be threatened globally (Nic Lughadha *et al*., 2020), and there is a risk that some may go extinct before being described (Roberts *et al*., 2016). Biodiversity hotspots, originally defined as regions containing exceptionally high concentrations of rare and endangered plant species, were conceptualised as a means to identify priority areas for conservation (Myers *et al*., 2000; Mittermeier *et al*., 2011). However, biodiversity hotspots and other more recent mapping efforts (e.g., Jung *et al*., 2021; Cai *et al*., 2023) have given relatively little consideration for taxonomic and geographic knowledge shortfalls. This in turn has the potential to misguide conservation actions, as newly described species are more likely to be threatened than others (Roberts *et al*., 2016), and a lack of species geographic data may result in both hotspot misidentifications (Meyer *et al*., 2015) and inaccurate estimates of extinction risk (Morais *et al*., 2013). While the use of an early version of WCVP has shown that biodiversity hotspots may be housing most undescribed plant species (Joppa *et al*., 2011), a thoroughly updated and joint consideration of taxonomic and geographic shortfalls could point towards new perspectives for collection and conservation.

Sampling effort is shaped indirectly by a wide array of environmental and socio-economic conditions (e.g., poverty, security, climate, topography, levels of environmental protection; Mair, 2016). These factors can sometimes represent insurmountable barriers to botanists, potentially making regions with the largest taxonomic and geographic knowledge gaps unrealistic targets for collection efforts, especially when considered alongside the relatively limited resources available to botanical organisations. However, regional socio-economic and environmental contexts are rarely considered when quantitatively assessing areas for collection (and conservation) prioritisation (but see Meyer *et al*., 2016). Accounting for regional opportunities and challenges is thus key to develop collection strategies that facilitate the description of the largest number of species, while also accounting for – and whenever possible helping to address – social and environmental issues of importance to civil society and botanists (Pascual *et al*., 2021).

Here, we pursue the challenge of accelerating the description of plant diversity by assessing taxonomic and geographic shortfalls, using WCVP and other databases such as the Global Biodiversity Information Facility (GBIF) as key data sources, and a suite of analytical methods. We specifically test (Q1) whether areas with the largest estimated taxonomic and geographic gaps overlap; (Q2) what attributes are most associated with the early description and geolocation of vascular plant species; and (Q3) whether global collection efforts of the previous decade have been concentrated in areas representing the largest expected shortfalls. Inspired by the concept of “dark taxa” (DNA sequences not identified to the species level; Page, 2016), we combine our estimates of taxonomic and geographic shortfalls to define plant diversity “darkspots” as areas housing most undescribed and poorly geolocated species, and assess (Q4) whether darkspots could become hotspots with increased taxonomic and geographic sampling efforts. Finally, we explore (Q5) a range of collection priority scenarios informed by a combination of key environmental and socio-economic drivers. Our results indicate substantial opportunities for revisiting plant collecting strategies to support the equitable conservation, sustainable use, and benefit sharing of the world’s plant diversity in line with the GBF.

## MATERIALS AND METHODS

We analysed historical and geographical patterns of description and geolocation of vascular plant species across *botanical countries* of the world, as defined by level 3 of the World Geographical Scheme for Recording Plant Distributions (WGSRPD; Brummitt, 2001). We used a time-to-event analysis (Moura & Jetz, 2021) to predict the number of vascular plant species remaining to be described and geolocated in each botanical country as a function of several species attributes (i.e., climatic and geographic ranges, uses by humans, and sampling effort). Then, we identified areas where both Linnaean and Wallacean shortfalls coincide (Q1); and species attributes driving these patterns (Q2).

Subsequently, we used a skyline modelling approach (Gjesfjeld, 2020) to compute past description and geolocation rates through time in each botanical country. We then tested whether our estimates of the potential number of undescribed and non-geolocated species are correlated with the collection rates of the previous decade across all botanical countries (Q3). Furthermore, we identified darkspots as areas containing the largest taxonomic and geographic shortfalls and assessed their (mis-)matches with known biodiversity hotspots (Q4). Finally, we characterised the socio-economic and environmental contexts of each botanical country using the United Nations’ Sustainable Development Goals (SDGs) indicators and a Principal Component Analysis (PCA). We then identified priority botanical countries for both the description of new species and the geolocation of known species under nine scenarios accounting for regional opportunities and challenges (Q5).

### Taxonomic and geographic data

WCVP (Govaerts *et al*., 2021) is a comprehensive global database that collates and reconciles information about most known vascular plant species from peer-reviewed literature, authoritative scientific databases, herbaria and observations; all of which is then curated by expert reviews. We used it to retrieve information on species nomenclature, synonymy, year of first global scientific description (*sensu* Linnaean taxonomy; DuBay *et al*., 2022), and geographic distribution at the botanical country level. Our study included only species with accepted names and excluded hybrids, resulting in a list of 343,523 vascular plant species. We obtained geo-referenced occurrence records for each of these species from the Global Biodiversity Information Facility (GBIF), the Botanical Information and Ecology Network (BIEN) (Enquist et al., 2016), BioTIME (Dornelas et al., 2018), NeoTropTree (http://prof.icb.ufmg.br/treeatlan) and speciesLink (Canhos et al., 2022, http://specieslink.net/). Data from GBIF was accessed through the Microsoft Planetary Computer (MPC) platform, a cloud computing environment with a catalogue of global spatio-temporal datasets (https://planetarycomputer.microsoft.com/; GBIF.org [01 March 2022] GBIF Occurrence Download, https://doi.org/10.15468/dl.nqbg5v).

We used exact-matching to link species names in GBIF’s taxonomic backbone to species in our WCVP list. We only kept names that matched directly with an accepted name or through homotypic synonymy. We standardised all metadata information using the DarwinCore format and used the *CoordinateCleaner* R package (Zizka *et al*., 2019) to exclude fossil occurrence records, records with common georeferencing mistakes, records with coordinates close to botanical institutions and GBIF headquarters, and duplicated records based on longitude, latitude, and year of collection. We removed all records outside each species’ native range according to WCVP. Our final dataset included >121 million occurrence records for 259,939 vascular plant species (c. 75% of our list of species retrieved from WCVP) across 450 families and 363 out of 369 botanical countries. For each species, we then extracted the year of collection of the first record found within each botanical country.

### Species-level explanatory variables

We considered ten variables that could affect the time to the description of a new species and the time to the geolocation of a species already described: annual mean temperature (AT), annual precipitation (AP), temperature seasonality (TS), precipitation seasonality (PS), elevation (E), range size (RS), taxonomic activity (TA), geographic activity (GA), number of uses by humans (USE) and life form (LF). For further details about the selection and compilation of these variables, please see Supporting Information (SI) Methods **S1a**.

To avoid collinearity issues in further modelling efforts, we detected highly correlated pairs of predictors (i.e., with a Pearson correlation coefficient higher than 0.7) and excluded the one of the pair with the greater Variance Inflation Factor (VIF; Zuur *et al*., 2010). As such, we excluded AT and all remaining predictors were then centred at their sample means and scaled by their sample standard deviations within their respective botanical country to help improve the inference of model parameters.

### Taxonomic and geographic shortfalls

We fitted two parametric hazard (Hz) regression models with two different response variables: (i) the number of years between 1753 (i.e., since Linnaeus’ first edition of Species Plantarum) and the description of species *i* (T_i,desc_), and (ii) the number of years between the description of species *i* and its first recorded occurrence point in botanical country *k* (T_k,i,geoloc_). For both models, we only considered species descriptions and geolocations up to the year 2021. We did not fit any model for Antarctica because it contained observations for only three species.

Since the year of geolocation is not available for all records, T_k,i,geoloc_ is considered *censored* (i.e., partially observed), which has been accounted for to allow for valid estimation of the model parameters (Hosmer *et al*., 2008) (see Supporting Information (SI) Methods **S1b**).

We used the Stan probabilistic programming language (Carpenter *et al*., 2017) through the *rstanarm* package (Goodrich *et al*., 2020) to fit these models in a Bayesian framework, using T_i,desc_ and T_k,i,geoloc_ as response variables, AP, TS, PS, E, RS, TA, GA, USE and LF as explanatory variables and the default Hamiltonian Monte-Carlo (HMC) sampler (Hoffman & Gelman, 2014) (for more details about the models’ parameterisation see SI Methods **S1c**). We ran two chains with 3,000 iterations, discarding the first 1,000 iterations as warm-up, leaving 2,000 (times 2) iterations as posterior samples. We assessed convergence using trace plots and the Gelman-Rubin diagnostic statistic (Gelman & Rubin, 1992), where values < 1.1 indicate adequate convergence.

The goodness of fit of both models was assessed using an integrated Brier score (iBS), which quantifies predictions accuracy over a given time period (see SI Methods **S1d**). iBS=0 indicates perfect accuracy of the model’s predictions, while iBS=1 indicates perfect inaccuracy.

We used the posterior samples of the models’ parameters to generate proportions of species remaining to be described and geolocated for each botanical country (for more detail about the procedure, see SI Methods **S1e**).

We used the predicted proportion of species remaining to be described to estimate the potential number of species awaiting scientific description as follows:

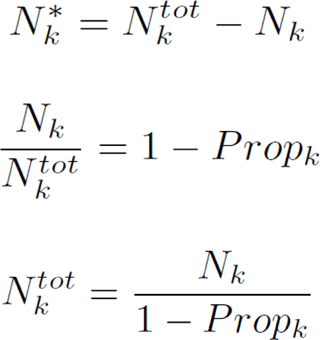

where, *N_k_\**denotes the number of species remaining unknown in botanical country k, *N_k_^tot^*and *N_k_* denote the total and currently known number of species in k, respectively, and *Prop_k_* denotes the aforementioned expected proportion.

To determine the potential number of species still awaiting geolocation, we calculated the number of species expected to occur in each botanical country according to WCVP, but with no valid occurrence records, then we multiplied this number by the expected proportion of species non-geolocated yet.

To facilitate comparison between botanical countries of different areas, we generated a second set of predictions rescaled to a fixed area of 10,000 km^2^ following (Brummitt & Lughadha, 2003).

### Effect of species characteristics

We extracted and exponentiated the regression coefficients of our Hz models from each botanical country to obtain hazard ratios (HRs) for each predictor variable. If HR=1, there is no effect of predictor variable *p*, if HR>1, an increase in *p* increases the chance of a species being described or geolocated, and if HR<1, an increase in *p* decreases this chance. Therefore, HRs allow to assess how and how much each species individual characteristic influences their description and geolocation.

### Past collection efforts

We used the software LiteRate (Koch *et al*., 2020) to fit a piecewise-constant model and jointly infer the rates of description and geolocation and the number and temporal placement of shift points (Silvestro *et al*., 2019; Gjesfjeld, 2020). The method uses a reversible-jump Markov chain Monte Carlo (rjMCMC) and its output can be used to obtain marginal rates through time that account for model uncertainty (Silvestro *et al*., 2019).

As inputs, we used the date of first scientific description and earliest geolocation in each botanical country. Species expected to be present in the botanical country according to WCVP but not found in our geographic occurrence data were considered not yet geolocated. We used a Poisson-death process to estimate the rates of species description (expected number of newly described species per year) and first geolocation within a botanical country (expected number of newly geolocated species per year among those already described).

We ran the analyses for each botanical country for 1.2 million rjMCMC iterations, sampling every 1,000 iterations. After assessing the convergence of the chains and excluding the first 20% of the samples as a burn-in, we averaged marginal rates across the period 2010-2019 (avoiding data collection patterns caused by the COVID-19 pandemic). Mean collection trends from 2010-2019 were then correlated with data gaps predicted for 2021 for both species description and geolocation using Kendall’s correlation in R package gpubr v.0.4.0.

### Darkspots and hotspots

To identify (mis-)matches between global biodiversity hotspots and the Linnaean and Wallacean shortfalls, we first obtained the geographic distribution of the 36 biodiversity hotspots from Hoffman (2016). To identify darkspots, we rescaled our Linnaean and Wallacean shortfalls estimates between 0 and 1, and summed their values to calculate a “darkspot score” ranging between 0 (i.e., smallest taxonomic and geographic knowledge gaps) and 2 (i.e., largest taxonomic and geographic knowledge gaps). To avoid arbitrary cut-offs and allow for a direct surface comparison, we categorised darkspots as botanical countries exhibiting the highest darkspot scores and a total cumulative land area equivalent to the total land area of the 36 biodiversity hotspots.

### Collection priorities

To characterise the average socio-economic development state of each botanical country, we used indicators monitoring progress towards all 17 SDGs from https://sdgs.un.org/goals. These relate to factors potentially impacting botanical expeditions, such as infrastructure and accessibility, security, access to vital resources, poverty, social inequalities, partnerships, levels of education and protection of the environment (Fig. S1). We assigned national values to botanical countries representing regions within countries.

The SDGs are all interlinked; therefore, we used a Principal Component Analysis (PCA) to summarise the variation in all SDG indicators across botanical countries. The first two Principal Components (PCs) explained 66.63% of the variance in the data (Fig. S1), and were used to determine our collection prioritisation scenarios. PC1 (hereafter, representing a proxy for “income group”) reflected variation in indicators associated with SDGs 1-13 and 16-17, and PC2 (hereafter, a proxy for “environmental protection”) was mainly associated with variation in SDGs 14 and 15 (Fig. S1).

We assume that botanical countries with high income and infrastructure will have a large potential to document their floras using their own resources. In contrast, prioritising botanical countries with lower income and a high potential for undescribed plant diversity can be used to highlight areas where the global community may need to join forces to support the country’s poverty alleviation, capacity building and funds, which are all likely required in order to achieve a full documentation of their floristic diversity. Similarly, prioritising botanical countries with low environmental protection can be used to highlight regions where species may have the potential of being lost if not described soon. Finally, prioritising botanical countries with high environmental protection may highlight areas where there may be more capacity and resources available for collecting plant data to inform and improve protected area management.

Our baseline scenario (Scenario 1) ranks botanical countries by the magnitude of the estimated taxonomic and geographic knowledge gaps only. We then defined Scenarios 2-9 using combinations between our baseline and the two extremes of the socioeconomic and environmental variables (i.e., prioritising collections based on income and/or environmental protection or not, and if so, prioritising botanical countries with more or less income and/or environmental protection). Each scenario is described in more detail in the Results section below.

To calculate the benefit of collecting in each botanical country under each scenario, we rescaled the darkspot score between 0 and 1, then summed with the income (PC1) and protection (PC2) scores of each botanical country rescaled between 0 and 1. We defined sets of global priorities for each scenario based on our darkspot definition. In addition, we defined continental priorities as the top five priority botanical countries by continent, as defined by the WGSRPD. Finally, to assess uncertainty in shaping priorities, we recalculated the benefit 10,000 times by resampling the mean and standard deviation of the estimated number of species to be described and geolocated. We then compared each resampling event with the main result to assess changes in the ranking of countries by using Spearman’s rank correlation.

## RESULTS

### Global distribution of the Linnaean and Wallacean shortfalls

Overall, our models performed well with Brier scores very close to zero indicating a good fit across all botanical countries (median iBS = 0.086, interquartile range = 0.017 for species descriptions; median iBS = 0.17, interquartile range = 0.058 for species geolocations).

Our results indicate that taxonomic and geographic knowledge gaps differ substantially in magnitude and vary widely across the globe (Table 1). The Linnaean shortfall is expected to be modest in most continental regions, with a vast majority of botanical countries predicted to contain less than 50 unknown species (Table 1). Yet, the Linnaean shortfall remains apparent in Asia-Tropical and South America (Fig. **1a** and Fig. **S2a**) with few botanical countries predicted to house hundreds of species still unknown to science (Table 1, Fig. **1a** and Fig. **S2a**), but also in Turkey, Madagascar and China south-central (Fig. **1a** and Fig. **S2a**). The Wallacean shortfall is predicted to be large in many parts of the world and exhibits a different global pattern, with thousands of known species still expected to be missing geographic records across most continents (Table 1, Fig. **1a** and Fig. **S2a**).

**Figure 1:**
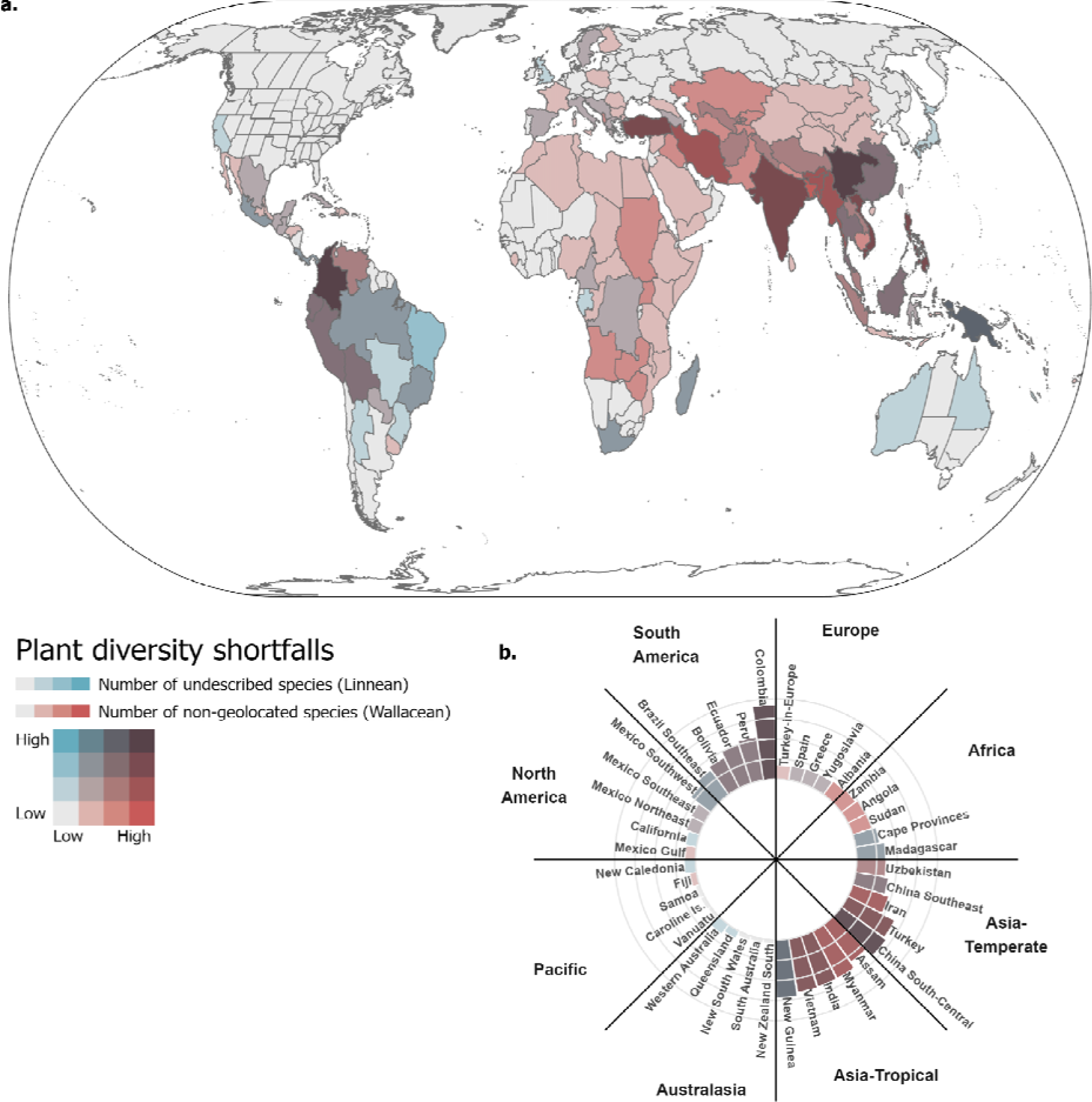
The global distribution of the Linnaean and Wallacean shortfalls for vascular plants. In panel **(a),** the Linnaean shortfall (blue gradient) is quantified as the predicted number of species yet to be described scientifically based on the estimated probability of species to remain undescribed by 2021. The Wallacean shortfall (red gradient) is quantified as the predicted number of species yet to be geolocated based on the estimated probability of species to have no geographic record available in a given botanical country by 2021. Colours indicate the relative magnitude of each shortfall, with light grey and dark brown colours representing small and large values for both shortfalls, respectively. In panel **(b),** the top 5 botanical countries with the largest combination of both Linnaean and Wallacean shortfalls are reported by continental region. Predictions were rescaled between 0 and 1 to allow for comparison.

**Table 1:** Summary of the predictions for the number of species remaining to be described (Linnaean shortfall) and geolocated (Wallacean shortfall) across all botanical countries of each continent.

| continent<sup>(1)</sup> | Number of undescribed species |  |  | Number of non-geolocated species |  |  | % botanical countries with x number of undescribed species<sup>(2)</sup> |  | % botanical countries with x number of non-geolocated species |  |
| --- | --- | --- | --- | --- | --- | --- | --- | --- | --- | --- |
|  | median | 2.5<sup>th</sup> percentile. | 97.5<sup>th</sup> percentile. | median | 2.5<sup>th</sup> percentile. | 97.5<sup>th</sup> percentile. | x≤50 | x>100 | x≤500 | x>1000 |
| Africa (N=71) | 9.0 | 3.0 | 63.8 | 346.0 | 35.2 | 1,886.1 | 95.8 | 1.4 | 58.6 | 24.3 |
| Asia-Temperate (N=52) | 8.0 | 2.3 | 119.3 | 509.5 | 87.9 | 3,605.1 | 92.3 | 3.8 | 50.0 | 26.9 |
| Asia-Tropical (N=30) | 19.0 | 3.7 | 153.9 | 1,913.0 | 60.9 | 4,512.1 | 60.0 | 16.7 | 16.7 | 66.7 |
| Australasia (N=13) | 13.0 | 2.6 | 52.7 | 70.0 | 22.6 | 249.3 | 92.3 | 0.0 | 100.0 | 0.0 |
| Europe (N=41) | 13.0 | 4.0 | 45.0 | 376.0 | 73.0 | 1,415.0 | 100.0 | 0.0 | 61.0 | 9.8 |
| North America (N=73) | 6.0 | 2.8 | 47.2 | 125.0 | 43.0 | 751.6 | 97.3 | 0.0 | 91.8 | 1.4 |
| Pacific (N=25) | 4.0 | 2.6 | 26.8 | 116.0 | 21.9 | 545.1 | 100.0 | 0.0 | 95.7 | 0.0 |
| South America (N=47) | 14.0 | 4.0 | 147.6 | 422.0 | 96.6 | 2,352.5 | 74.5 | 12.8 | 57.4 | 12.8 |

<sup>(1)</sup>N is the number of botanical countries included.
<sup>(2)</sup>% refers to the proportion in percent.

Overall, Colombia, New Guinea, China south-central, Vietnam, and India have the greatest combined Linnaean and Wallacean shortfalls globally, in decreasing order (Fig. 1). By continent, New Caledonia and Fiji have the greatest combined shortfall for the Pacific, Western Australia and Queensland for Australasia, New Guinea and Vietnam for Asia -Tropical, China South-Central and Turkey for Asia - Temperate, Madagascar and Cape Provinces for Africa, Albania and Yugoslavia for Europe, Mexico Southwest and Mexico Southeast for North America, and Colombia and Peru for South America.

### Species attributes associated with early description and geolocation

Assessing the importance of different species attributes on their early description and geolocation, we find that range size is one of the most important drivers of the Linnaean shortfall: largely distributed species are consistently more likely to be described before small-range species, which also indicates that recently described species tend to be characterised by smaller ranges (Fig. 2). Conversely, species that are rapidly geolocated are more likely to exhibit small geographic ranges, while species that are geolocated more slowly are more likely to have large geographic ranges.

**Figure 2:**
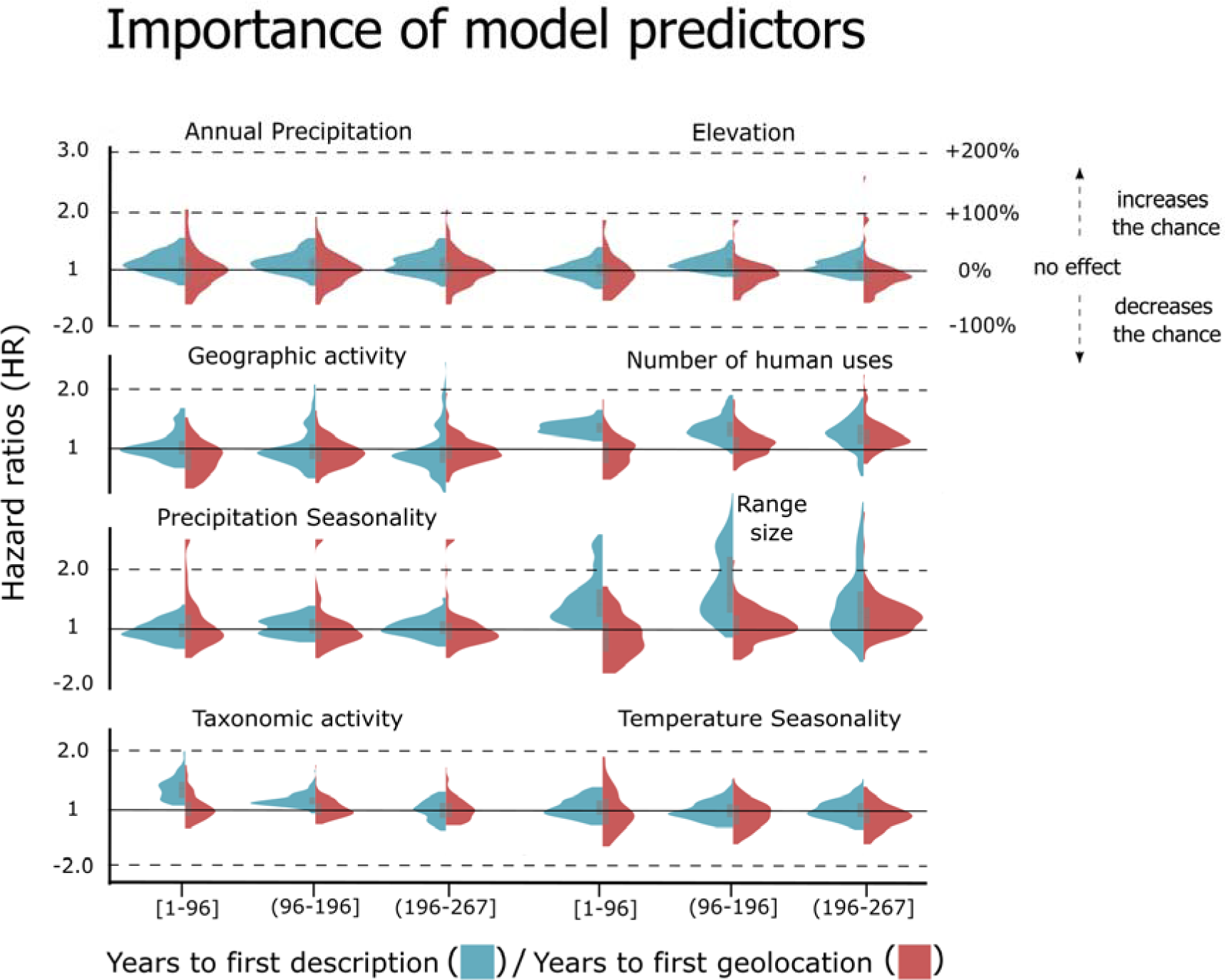
Importance of variables for the description and geolocation of vascular plant species. The violin and box plots show the 95% posterior distribution of the hazard ratio (HR) (exponential of the time-to-event model coefficients). HR measures the change in the chance of describing or geolocating a new species in a given botanical country following an increase in the considered variable. For example, if a time-to-description model has one variable, such as range size, with a HR of 1.5, it means that a species with a range covering three botanical countries has |1 - 1.5| x 100 = 50% more chance to be described than a species with a range size of two. In other words, the time to describe a species with a range size of three is reduced by 50% compared to a species with a range size of two, meaning that it would take twice as long to describe a species with a range size one-unit smaller.

We find that species with a high number of uses by humans are more likely to be described earlier than species with fewer uses (Fig. 2). Uses by humans are also associated with accelerated geolocation, but mostly in species geolocated belatedly. Early descriptions are positively related with taxonomic activity at a given time, while no clear pattern emerges for species geolocation. The magnitude and direction of the effects of geographic activity, elevation, temperature seasonality and annual precipitation strongly vary across botanical countries such that none of these environmental variables seem to consistently influence species description or geolocation in one direction or another.

### Past collection dynamics and plant diversity shortfalls

Species descriptions first peaked in Europe and large parts of North America and Australasia in the 1850s, before peaking in most of Africa, Asia-Temperate and the Pacific in the last century (Fig. **S3a** and **S3b**). Species geolocations peaked in eastern Russia and North Africa in the 1850s, and more recently in Europe, South East Asia and Sub-Saharan Africa (Fig. **S3c** and **S3d**).

We find a mismatch between the geographic distribution of collection rates over the 2010-2019 period and the distribution of gaps. Kendall’s correlations tests highlighted weakly positive but significant correlation between description rates in 2010-2019 and the predicted number of species remaining to be described (Fig. **3a** and Fig. **S4a**). We also find a stronger and significant correlation between geolocation rates (2010-2019) and where most species are predicted to remain to be geolocated in the future (Fig. **3b** and Fig. **S4b**).

**Fig. 3:**
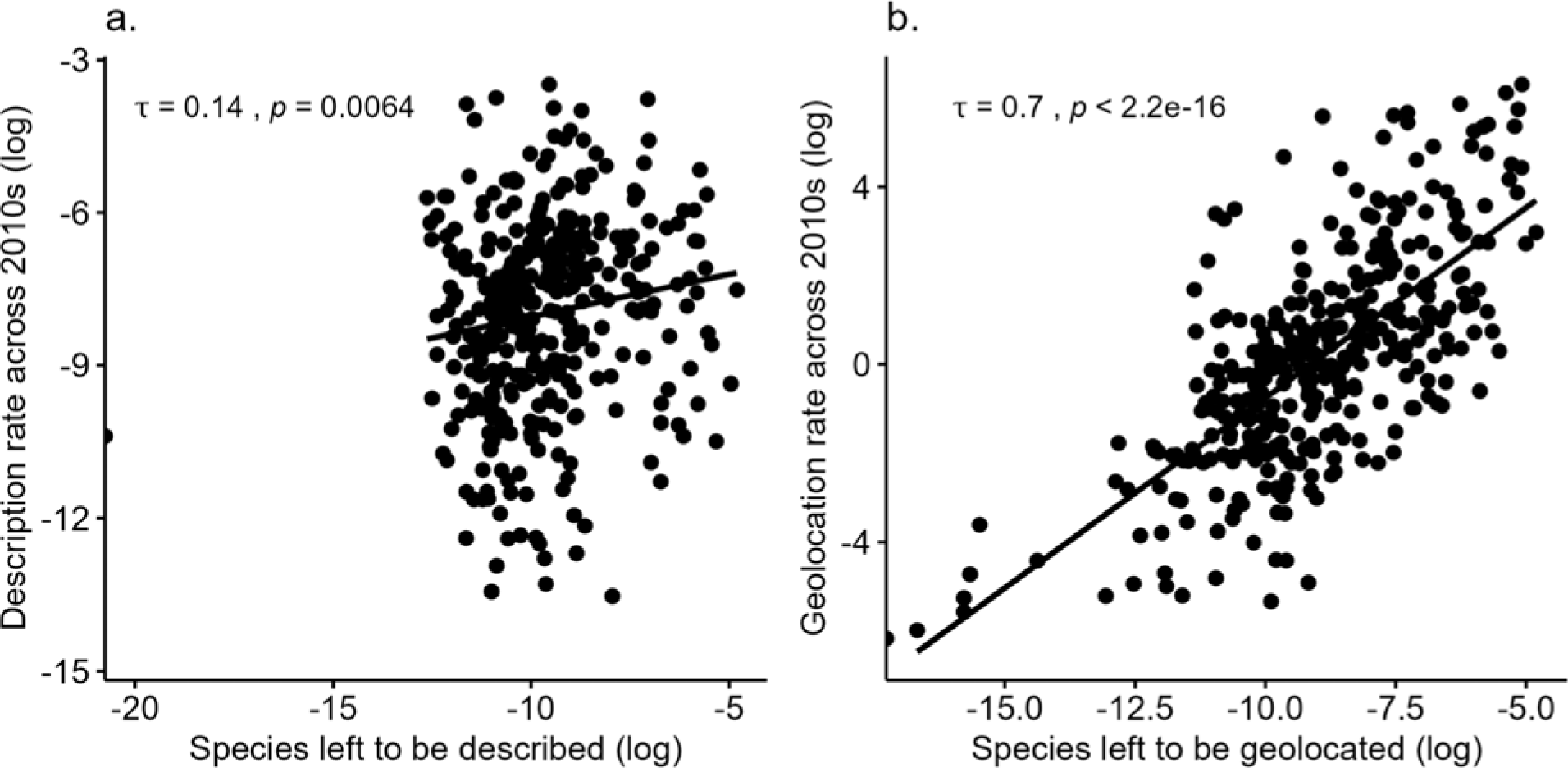
Kendall’s correlations between past collections and future needs. Dots represent botanical countries. In panel **(a),** the x-axis represents the log-transformed predicted number of species remaining to be described and the y-axis represents the log-transformed average species description rate estimated across 2010s. The weak positive relationship means there is a slight trend of higher description rates in botanical countries that are predicted to have more species remaining to be described. In panel **(b),** the x-axis represents the log-transformed predicted number of species remaining to be geolocated and the y-axis represents the log-transformed average species geolocation rate estimated across 2010s. The positive relationship means sampling has recently occurred in areas that are predicted to have more species remaining to be geolocated. Recent sampling has thus been more effective in bridging the Wallacean than the Linnaean shortfall.

### Plant diversity darkspots and biodiversity hotspots

Vascular plant diversity darkspots are represented by botanical countries exhibiting the highest darkspot scores globally (i.e., sum of the estimates of the rescaled Linnaean and Wallacean shortfalls), and a cumulative area equal to that of biodiversity hotspots to allow for comparison. We identified 32 darkspots: 14 across large parts of Asia - Tropical (Myanmar, Assam, Philippines, Vietnam, Bangladesh, New Guinea, India, East and West Himalaya, Thailand, Sumatera, Laos, Malaya, and Borneo), nine in South America (Colombia, Peru, Venezuela, Brazil North, Brazil Southeast, Ecuador, Costa Rica, and Panama), six in Asia - Temperate (China South-Central, Turkey, Iran, Uzbekistan, China Southeast, and Tadzhikistan), two in Africa (Madagascar and Cape Provinces), and one in North America (Mexico Southwest).

We find that the 32 darkspots overlap to a very large degree with the 36 biodiversity hotspots, with the notable exception of New Guinea (Fig. **4**). In contrast, there is a large number of hotspots that are not identified as darkspots in our analysis, particularly in the Pacific, Australasia, North America, Africa, and Europe. When re-scaling the darkspot score to a fixed area, 27 new botanical countries (e.g., Haiti, Zambia, Albania, Afghanistan, Nepal and Cambodia) become darkspots (Fig. **S5**).

**Figure 4:**
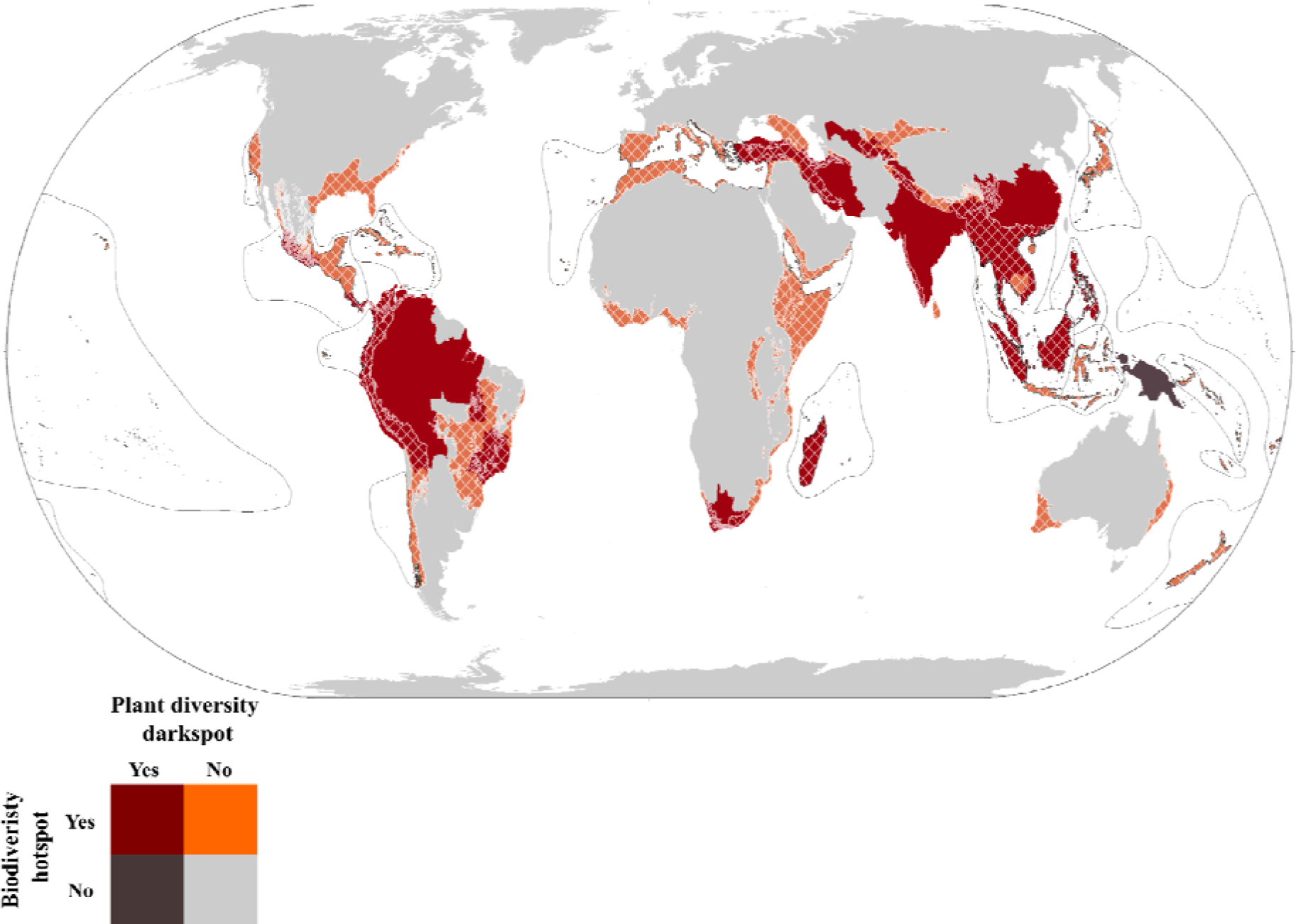
Global darkspots of vascular plant diversity and biodiversity hotspots. Dark red indicates botanical countries that are identified as darkspots and that contain hotspots. Black indicates botanical countries uniquely defined as darkspots. Orange indicates areas containing hotspots but not overlapping with any darkspot, and grey indicates areas containing neither darkspots nor hotspots. The cross-hatched areas and black lines indicate the terrestrial and marine delimitation of biodiversity hotspots, respectively. All darkspots overlap with hotspots, except New Guinea.

### Plant collection priorities

We defined nine scenarios based on different trade-offs between the darkspot score, the income group, and level of environmental protection (Fig. **5** and Fig. **S6**). Scenario 1 is considered our baseline scenario and is represented by the darkspot score with no consideration for the botanical countries’ level of income or environmental protection (Fig. **1** and Fig. **S2**). Scenarios 2-9 highlight regional and global collection priorities deviating from the baseline and varying greatly across space according to different income-protection combinations (Fig. **5**, Fig. **S6-S14**). However, consistent global priorities arise across all scenarios: Borneo, Colombia, Ecuador, India, Myanmar, New Guinea, Peru, the Philippines and Turkey (Fig. **5**). When re-scaling the darkspot score to area, India is no longer a global priority across all scenarios, while Assam, Bolivia, Costa Rica, Cape Provinces, East Himalaya, Laos, Maldives, Malaya, Mexico Southwest, Panama, Selvagens, Thailand, Uzbekistan and Vietnam become consistent global collection priorities (Fig. **S6**).

**Figure 5:**
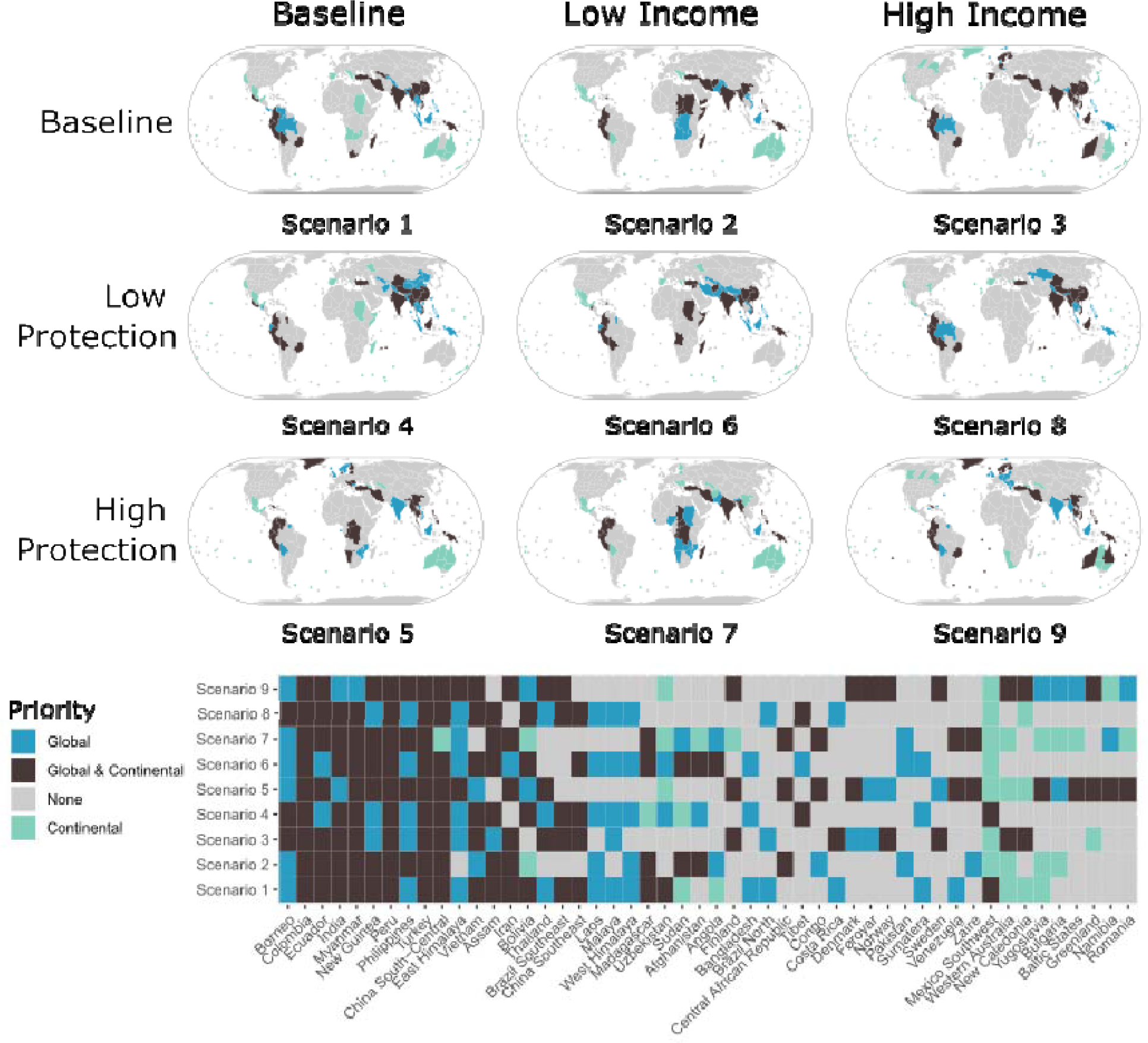
Collection priorities under nine income and environmental protection scenarios. Different scenarios investigate different tradeoffs between the darkspot score (i.e., sum of scaled taxonomic and geographic shortfalls), income group (i.e., first axis of the Principal Component Analysis (PCA) summarising the variation of the Sustainable Development Goals’ indicators across botanical countries) and environmental protection level (i.e., second axis of the PCA). Top global (blue) and continental (green) collection priorities are provided across scenarios, as well as their overlap (black) and absence (grey). The bottom matrix provides more detail on top priority countries across the nine scenarios. Borneo, Colombia, Ecuador, India, Myanmar, New Guinea, Peru, the Philippines and Turkey appear consistently as global priorities across all scenarios, providing high potential for new species descriptions and geolocations. Collections in other botanical countries will depend on tradeoffs between income and environmental protection.

When prioritising botanical countries with low income levels (i.e., Scenarios 2, 6 and 7), Afghanistan, Pakistan, Sudan, and Angola become global priorities consistently, while Thailand, Brazil Southeast, Brazil North, Mexico Southwest and Costa Rica are not considered global priorities anymore (Fig. **5**, Fig. **S6-S14**). Conversely, when prioritising high-income botanical countries (i.e., Scenarios 3, 8 and 9), no botanical country becomes a global priority consistently across all scenarios, while Madagascar, Uzbekistan, Bangladesh, Sumatera, Venezuela, and Mexico Southwest are consistently removed from the global priority list (Fig. **5**, Fig. **S6-S14**).

When prioritising botanical countries with low levels of environmental protection (i.e., Scenarios 4, 6 and 8), Tibet becomes a global priority consistently, while Venezuela is consistently removed from the global priority list (Fig. **5**, Fig. **S6-S14**). Finally, when prioritising botanical countries with high levels of environmental protection (i.e., Scenarios 5, 7 and 9), no botanical country becomes a global priority consistently, while China Southeast, Laos, West Himalaya, Malaya, Uzbekistan, Bangladesh, Brazil North, Sumatera, Costa Rica, and Mexico Southwest are no longer considered global priorities (Fig. **5**, Fig. **S6-S14**). Finally, we found that the average rank of randomised runs was generally consistent with our prioritisation (Figs. **S15-S23**, Spearman’s rank correlation 0.9853-0.9988, p< 2.2e-16), except for New Guinea, Borneo, Ecuador, Bolivia, Brazil North and Panama, which increased in priority and could be darker spots than first predicted.

## DISCUSSION

### Global distribution of the Linnaean and Wallacean shortfalls

Using the most extensive dataset to date on standardised plant distributions at the botanical country level, our research provides a joint assessment of the global taxonomic and geographic data shortfalls. Our predictions indicate that a complete documentation of the geographic distribution of all vascular plants will be slow to achieve if botanical collections continue to follow past trends of species description and geolocation (Fig. 3). An incomplete understanding of plant diversity and occurrences could jeopardise actions towards implementation of some of the goals of the GBF, including protecting and restoring plant diversity.

Predicting numbers of species remaining to be documented comes with high uncertainty (Bebber *et al*., 2007; Moura & Jetz, 2021). Although the models developed here performed well, confidence intervals around our predicted absolute numbers are inevitably large (Table 1). Even so, relative comparisons across botanical countries of the world reveal well-supported insights. Many species are predicted to remain undescribed in areas of the world already known for their exceptional levels of plant endemism (Kier *et al*., 2009), generally suggesting that there may be more species to describe where there are more narrowly distributed species currently known to science. Expected gaps in geographic records follow very different distribution patterns, with larger numbers of species remaining to be geolocated in both species-rich and species-poor regions, mainly in eastern regions from North Africa to the Indo-Burma region, Central Asia and the Middle East.

Our results suggest that the Wallacean shortfall may be driven by other factors than the intrinsic characteristics of plant diversity; in particular, they may be more related to political, economic, or socio-cultural conditions, as highlighted in previous publications (Fig. 2, Figs. S3-4; Meyer *et al*., 2015, 2016; Zizka *et al*., 2021). Filling both knowledge gaps will therefore imply actions at global to local scales, from policy, finance and science, to ultimately influence where future botanical surveys should be conducted.

### Species-level drivers of plant description and geolocation

Our results corroborate studies conducted on other biodiversity data shortfalls (Rudbeck *et al*., 2022; Maitner *et al*., 2022), by showing that species range size is a major driver of knowledge gaps (Hortal *et al*., 2015). Similarly to vertebrates (Essl *et al*., 2013; Moura & Jetz, 2021), plant species with large geographic ranges are described earlier than small-ranged species (Fig. 2). Since they are also more likely to be rare locally (Brown, 1984), narrowly-distributed species may be harder to detect and thus remain scientifically unknown.

Conversely, the geolocation of widespread species takes considerably longer, which may be explained by the fact that more ground needs to be covered to record their geographic position, and that their commonness makes them less sought-after by botanists (Fig. 2; Daru *et al*., 2018). Additionally, the higher the socio-eco-cultural value of a species (as measured by the number of uses by humans), the sooner it is likely to be studied (Adamo *et al*., 2021), described, and to a lesser extent geolocated. Species described longer ago may also have had more time to be used in a variety of ways across the world. Species’ appeal to botanists generally accelerates their description, while it does not relate to their geolocation. This follows the documented pattern that botanists focus more on providing first observations of as many species as possible than on providing multiple observations of the same species across its geographic range (Steege *et al*., 2011). Aligning both taxonomic and geographic sampling efforts will be key to filling plant diversity shortfalls. Taken together, our results suggest that most currently unknown plant species may be from groups of taxa neglected for taxonomic and societal reasons, occupying restricted ranges across diverse environmental conditions.

### (Mis-)matches between past trends and future needs

Our findings align with previous studies highlighting spatial and temporal biases in species collections shaped by the history of botanical sciences, the advent of technologies (e.g., Global Positioning Systems), geographic patterns of species richness and other socio-economic factors (Fig. **S3**; Meyer *et al*., 2015, 2016; Farooq *et al.,* 2021; Hughes *et al*., 2021).

Our analyses also feature a positive overall correlation between where collections have happened in the recent past and where the largest data gaps are estimated to remain in the future, which indicates a good general progress towards bridging the Linnaean and Wallacean shortfalls (Fig. **3** and Fig. **S4**). However, while the correlation is relatively strong for geolocations, it is weaker for species descriptions. We interpret this difference with caution given that predicting description needs may come with higher uncertainties than for geolocations. In fact, predictions of potentially missing geolocations are based on expected targets defined by the known species richness of each botanical country, whereas predictions of future gaps in descriptions cannot be informed in such a way. In addition, it is important to note that predictions of gaps in geolocation do not account for currently undescribed species. Beyond uncertainty, large recent description rates across islands (e.g., Kriti, East Aegean islands, Mozambique Channel islands, Comoros) may indicate preferences for areas with high endemicity levels and constrained spaces to search for new species (Fig. **S3b;** Kier *et al*., 2009). Botanical countries such as New Zealand, British Columbia and Ireland have attracted extensive description efforts recently, but have limited predicted numbers of species remaining to be described in the future. Conversely, several areas containing potentially large numbers of undescribed species (e.g., Peru, Colombia and New Guinea) have had relatively less attention and would require immediate strategic investments to expand botanical science capacity.

### Knowledge gaps and conservation priorities

By identifying darkspots of vascular plant diversity and comparing their distribution to that of the biodiversity hotspots, we intended to assess whether some regions could represent neglected conservation priorities globally. Our results follow previous assessments based on estimates of the Linnaean shortfall, indicating a general overlap between collection and conservation priorities (Joppa *et al*., 2011).

New Guinea, the island with the largest number of plant species (Cámara-Leret *et al*., 2020), is the only identified darkspot that is not considered a hotspot (Fig. 4; Joppa *et al*., 2011). Under the most recent hotspot assessment, New Guinea was recognized as an extremely rich region in endemic species but it did not meet the criterion of having 30% or less of its original natural vegetation left intact and thus was not considered to be threatened enough (Mittermeier *et al*., 2011). Given land-use changes primarily associated with agriculture are increasingly threatening those regions (Brun *et al*., 2015), New Guinea may require particular attention before many of its known and yet-to-be-known species become threatened, or possibly extinct. The Brazil-North darkspot overlaps slightly with the Tropical Andes and Cerrado biodiversity hotspots, although it mostly comprises the Amazonian rainforest - a species-rich region that still conserves a large fraction of its native vegetation, while experiencing increasing threats (Lapola et al., 2022). Thus, Brazil-North and the Amazonian rainforest may be in a similar position as New Guinea and require special conservation attention despite not being considered a biodiversity hotspot.

Beyond the hotspot classification (which has a range of limitations; Mace *et al*., 2000), considering plant diversity darkspots could have wider implications for the identification of priority areas and species for conservation (e.g., Jung *et al*., 2021), including for the International Union for Conservation of Nature’s Red List of Threatened Species (Rodrigues *et al*., 2006), and the designation of Key Biodiversity Areas (KBAs) and (Tropical) Important Plant Areas ((T)IPAs; Darbyshire *et al*., 2017).

### Future collection priorities

There are many nuances involved in how, where, when and why specimens are collected (Meyer *et al*., 2015). Yet despite considering a wide range of prioritisation scenarios accounting for different socioeconomic and environmental trade-offs (Fig. 5 and Fig. S6), some botanical countries are consistently identified as global priorities: Borneo, Colombia, Ecuador, India, Myanmar, New Guinea, Peru, the Philippines and Turkey. Amongst these botanical countries, some are predicted to contain relatively large numbers of species remaining to be described (e.g., New Guinea), geolocated (e.g., Myanmar), or both (e.g. Colombia), and all overlap with biodiversity hotspots.

Our results add to a body of literature showing that decision-making can be robust to uncertainty (McCarthy *et al*., 2003; McDonald-Madden *et al*., 2011), with clear win-win sampling strategies across scenarios. However, decision uncertainty increases as darkspots become less strong and tradeoffs between socioeconomic and environmental values come into play to drive the prioritisation. Our results (especially Fig. 5) provides an evidence-based and flexible framework to prioritise future collection efforts, which highlights the importance of a coordinated and inclusive effort towards tackling plant diversity shortfalls globally.

Data portals have changed the face of biodiversity science in recent decades and are still expected to play a major role in the future (Nelson & Ellis, 2019). Despite being one of the most complete data portals for species observation, GBIF does not include information from several existing (sub-)national databases. This has likely impacted our findings by suggesting some spots are “darker” than they truly are. It will therefore be key to integrate and share species information coming from these existing databases, as well as other collections (Heberling *et al*., 2021). In fact, the digitization of some of the largest herbarium collections in the world has contributed to filling major knowledge gaps, but this effort is largely incomplete and remains a priority (Nelson & Ellis, 2019). Moreover, many smaller-scale (and often under-resourced) herbaria containing exceptionally unique information remain inaccessible and vulnerable globally (Paton *et al*., 2020).

Citizen science can also greatly improve our botanical knowledge, as it can require less resources, it relies greatly on individual decisions, and allows for the upscaling of science outreach and botanical education (Pocock *et al*., 2018; Callaghan, 2022). Many identified darkspots contain large human populations and thus many potential contributors to citizen science projects (Cincotta *et al*., 2000). On the other hand, vouchered botanical specimens remain essential to a large number of scientific projects, and can seldom be replaced by photographs or undocumented observations (Troudet *et al.,* 2018).

Implementation of the actions described above has several fundamental requirements (that are also potential barriers). These include supporting botanical sciences and institutions financially, and enhancing the education and development of a new and diverse generation of plant taxonomists – both of which will be key to fill knowledge gaps with high-quality information (Paton *et al*., 2020). It is fundamental to incentivize data collection, integration, and sharing, not only in global policy frameworks but also at (sub-)national scale. In this context, our analyses may have limited reach given their coarse resolution, and much effort will be required in the future to provide assessments of data collection (and subsequently conservation) priorities at higher spatial resolution within countries. General levels of development and botanical sampling are also intertwined: access to food, water, medicine, and education, and the presence and quality of general infrastructure to travel, work and live safely are essential factors required to support the successful conduct of botanical activities (Meyer *et al*., 2015; Paton *et al*., 2020).

While using botanical information is necessary to develop efficient strategies to conserve plant diversity, delaying the protection of species and areas may lead to the extinction of species before they are even named scientifically and geolocated. The collection and preservation of global plant diversity is urgent, for their intrinsic value, to safeguard ecosystems, and to enhance people’s awareness and well-being (Antonelli *et al*., 2020; Pironon *et al*., 2020).

## ACKNOWLEDGEMENTS

We thank Steven Bachman for suggesting to expand the concept of “dark taxa” to darkspots of plant diversity, and Rhian Smith and Marybel Soto Gomez for edits on the final version of the manuscript.

## AUTHOR CONTRIBUTIONS

Samuel Pironon (SP) and Alexandre Antonelli (AA) conceived the paper with support from Ian Ondo (IO), Kiran Dhanjal-Adams (KDA), and Daniele Silvestro (DS). IO and Matheus Colli-Silva (MCS) compiled and processed spatial datasets, with support from Nicky Nicolson (NN) and Barnaby Walker (BW). KDA compiled the SDG datasets. IO, KDA, SP, and DS contributed to the analyses with support from AA. IO performed the time-to-event analyses, KDA performed collection trend analyses with support from SP, and DS performed the skyline modelling. IO, KDA and SP designed and produced the figures. IO, KDA, and SP led the writing with support from DS and AA. BW, DS, Alexandre K. Monro (AKM), Victor Deklerck (VD), NN and Olwen Grace (OG) provided feedback during the interpretation of results. All authors revised and approved the final version of the manuscript.

## SUPPORTING INFORMATION

All analysis was performed in R (version 4.1.3), except retrieving data from MPC and modelling past collection patterns, which were performed in python (version 3.10). We used the python package *pystac* to search the MPC catalogue and downloaded the resulting data using the *planetary-computer* package.

## DATA AND CODE AVAILABILITY

The data and code that support the reproduction of the findings and figures of this study will be made openly available at [repository name] and https://github.com/DarkSpots respectively, upon publication.

The LiteRate program is available here: https://github.com/dsilvestro/LiteRate and the version used in this manuscript is included in the [repository name].

