## Supplementary material for "Plant diversity darkspots for global collection priorities": Methods S1

**Methods S1** Section (a) describes how we selected and obtained species, (b) describes how we handled missing geolocation information, (c) how we parametrised our time-to-event models, (d) how we assessed the calibration of our time-to-event models and (e) how we made predictions for each botanical country.

**a. Selecting and compiling species-level covariates for the time-to-event analysis.**

The first five abiotic variables mean annual temperature (AT), annual precipitation (AP), temperature seasonality (TS), precipitation seasonality (PS) and elevation (E), were shown to impact the time to description of a range of tetrapod species (Moura & Jetz, 2021), and could likely also influence vascular plant descriptions. That study hypothesised that many western naturalists and large collectors had been initially trained at low elevation and in temperate environments, which may have shaped taxonomic and geographic collections through time. We obtained raster grids at 1km resolution for AT, AP, PS, and TS from the Climatologies at High resolution for the Earth’s land Surface Areas (CHELSA; Karger *et al.*, 2016), and for E from the Global Multi-resolution Terrain Elevation Database (Danielson & Gesch, 2011). We upscaled these layers to a ~110km resolution (c. 1 decimal degree). For each of the 249,521 species with occurrence records, we calculated the mean value of each variable over grid cells occupied by those occurrences. For the 94,233 species with no occurrence record, we averaged variables across the native botanical countries of each species.

Range size is also known to influence the time to species description since widespread species are often more likely to be detected earlier than narrowly distributed species (Enquist *et al.*, 2019; Moura & Jetz, 2021). We calculated range size as the total area of botanical countries comprising the species’ native range according to the WCVP. Taxonomic and geographic sampling is known to be biased towards certain taxonomic groups due to collector preference (Moura & Jetz, 2021), or because species fulfil particular needs for human well-being, such as food and medicine (Lucena *et al.*, 2012). Thus, we computed GA as the number of occurrence records collected within the native range of a focal species, divided by the number of years between the earliest and the latest (first) record collected within this range. We computed TA following Moura & Jetz (2021) as the number of named authors in the WCVP describing species from the same family during the year of description of the focal species, divided by the number of species described within the given family that year. To estimate USE, we obtained information about the uses of plant species by people for ten use categories (i.e., human food, vertebrate food, invertebrate food, medicine, poison, material, fuel, environmental uses, gene sources, and social uses) from the World Checklist of Useful Plant Species (Diazgranados *et al.*, 2020). We counted the number of use categories reported for each species to obtain a coarse estimate of the utility value given to species by humans. We categorised the life form (LF) information from the WCVP database into 4 classes following Humphreys et al. (2019): annual, epiphyte, herbaceous perennial and woody perennial.

We imputed missing life forms for 86,807 species (ca 25% of the total data) using the R package missForest (Stekhoven, 2022) which implements a non-parametric machine learning imputation approach based on random forests (Stekhoven & Buehlmann, 2012). We ran the missForest algorithm with 100 trees, using AT, AP, PS, TS, E, and RS as log-transformed predictors to improve the performance of the algorithm. We further included the family name of the species to improve the accuracy of the imputation.

**b. Censoring of the time to the geolocation of a species.**

We distinguish species with a GBIF record in a botanical country that lacks a year of collection (left-censoring) and species that are recorded in a botanical country by the WCVP with no GBIF record in that country (right-censoring). Where no year of geolocation was available, we used the time between description and 2021 as *T_k,i,geoloc_*. Therefore, *T_k,i,geoloc_* represents the maximum time-to-event for the left-censored records and the minimum time-to-event for the right-censored records. We therefore observed:

$$T_{k,i,geoloc}=\{T_{k,i,geoloc}^{*}, if no censoring. max\left( T_{k,i,geoloc}^{*},C_{k,i,geoloc} \right), if left-censoring. min\left( T_{k,i,geoloc}^{*},C_{k,i,geoloc} \right), if right-censoring.$$

where *T* ${}^{*}$*_k,i,geoloc_* is the ‘true’ time event and *C_k,i,geoloc_* is the censored time event, calculated as the number of years between the description of species i and 2021.

We defined an event indicator variable *d_i,k_*, which indicates whether *T_k,i,geoloc_*s were exact times, and left-censored or right-censored times. This allow to create an object of class “Surv” with the function *Surv* from the R package **survival** (Terry M. Therneau & Patricia M. Grambsch, 2000), which helps to code the different censoring cases just described. The model is then fitted with the function *stan_surv* from the R package **rstanarm** (Goodrich *et al.*, 2020), using this object as response variable.

**c. Parametrisation of the hazard models of the time to species description and geolocation.**

Our models define a baseline chance of an event occurring over time as a hazard function $h_{0}\left( t \right)$ scaled by the set of predictor variables $\eta_{ij}$, namely annual precipitation, temperature and precipitation seasonality, elevation, range size area, taxonomic activity, geographic activity, lifeform and number of uses by humans. We used the function coxph and cox.zph from the survival package to run preliminary models and test whether or not to include time-varying or time-fixed effects, i.e. treating the effects of variables as varying or not as a function of time, which is equivalent to assuming non-proportional or proportional hazards respectively. Based on these runs, only continuous variable effects (i.e. all but the lifeform) were allowed to vary with time when significant time-varying effects were detected. We added a random effect on the plant family (also known as a ‘frailty’ term) to account for potential correlations that could arise if species in the same family shared similar chances of being described and geolocated as a result of collector biases. Therefore, our models take the form:

$$h_{i}\left( t|X=x \right)=h_{0}\left( t \right)*exp\left( \eta_{ij} \right) h_{0}\left( t \right)=\sum_{l=1}^{L} \gamma_{l}M_{l}\left( t;k,\delta\right), \delta=3 , k=\{k_{1},...,k_{10}\} \eta_{ij}=\beta_{0}+x_{i}^{T}\beta_{p}+b_{j}^{T}Z_{ij}$$

where, *h_i_(t|X=x)* denotes the hazard rate for the $i^{th}$ species at time *t* given the vector of individual attributes *x_i_*, *h_0_(t)* denotes the baseline hazard rate, $\beta_{0}$ denotes an intercept parameter, $\beta_{p}$ denotes the vector of *P* coefficients of each covariate, *Z_i,j_* denotes the vector of covariates of the $i^{th}$ species in the $j^{th}$ plant family, *b_j_*, denotes the family-specific (random effect) parameter, $\gamma_{l}$ denotes the $l^{th}$ M-splines coefficient, and *M_l_* denotes the $l^{th}$ basis term for a degree $\delta=3$ (cubic) M-splines function evaluated at the vector of knot locations *k*.

The quantity exp($\beta_{p}$) refers to the *hazard ratio* (HR), which measures the relative increase in the hazard associated with a one-unit increase in the predictor *p*. If HR = 1, there is no effect of *p*, if HR > 1, an increase in *p* increases the hazard, and if HR < 1, an increase *p* decreases the hazard.

We assigned a normal prior distribution with mean 0 and standard deviation of 20 for $\beta_{0}$, and a standard deviation of 2.5 for $\beta_{p}=\left( p=1,...P=7 \right)$, and assigned a Dirichlet prior with a concentration parameter vector of ones for the M-splines coefficients:

$$\gamma_{l=1,...L}\sim Dirichlet_{L}\left( 1,1,1,... \right) \beta_{0}\sim N\left( 0 , 20 \right) \beta_{p=1,...P}\sim N\left( 0 , 2.5 \right) b_{j}\sim N\left( 0 , \Sigma_{b} \right)$$

where, $\Sigma_{b}$ is a variance-covariance matrix.

We used cubic B-splines for time-varying effects $\beta_{p}$(t) such that:

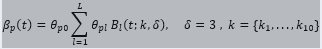

where 𝜃*_p0_* is a constant, *B_l_(t; k,* 𝛿*)* is the *l^th^* *(l = 1,...,L)* basis term for a degree 𝛿 B-spline function evaluated at a vector of knot locations *k = {k*_1_, …, *k*_J_*}*, and 𝜃*_pl_* is the *l^th^* B-spline coefficient.

**d. Evaluating models fit using the integrated Brier score (iBS).**

We generated 2,000 posterior predictions (using *posterior_survfit* as in section **(e)**) and used the functions *pec* and *crps* from the R package **pec** (Mogensen *et al.*, 2012) to first calculate the Brier score (BS) for each unique time we want predictions for (default in *posterior_survfit* is to use all unique times observed in the data). BS represents the averaged square difference between the predicted probabilities and the event indicator variable *d_i,k_* (described in section **(b)**).

In absence of censoring, BS at a given time t is calculated as:

$$BS\left( t \right)=\frac{1}{n_{k}}\sum_{i=1}^{n_{k}} \left( 1_{T_{i}>t}-S_{i,k}\left( t \right) \right)^{2}$$

where, *S_i,k_(t)* denotes the predicted probability for species i in botanical country k at time t, $1_{T_{i}>t}$ is the event indicator *d_i,k_* and *n_k_* denotes the number of species used for model fitting in k.

If the dataset contains censored observation, then BS(t) is adjusted by weighting the squared distances using the inverse probability of a censoring weights (IPCW) method, that is the “marginal” censoring model (Kaplan & Meier, 1958). Then, we computed the iBS using the function *crps* that integrates BS values over the range of times specified earlier, and we averaged the score for each botanical country. The function *pec* can only account for right-censored observations, so we calculated iBS for left-censored observations using the interval-censored implementation used by the function *sbrier_IC* from the **ICcforest** package (Yao *et al.*, 2020).

**e. Generating standardised predictions by botanical country.**

We used the function *posterior_survfit* from the R package **rstanarm** with the argument “type” set to “surv” to indicate that we want to predict survival probabilities, and the argument “times” set to different time spans depending on the model:

- To generate 2,000 posterior predictions for the probability that a species would remain undescribed, we set “times” to the number of years between 1754 and the year of the most recent record up to 2021.
- To generate 2,000 posterior predictions that a species would remain non-geolocated, we filtered species missing geolocation and set “times” to their *T_k,i,geoloc_* (cf. section **(a)** and the methods in the main manuscript).

We then averaged these species-level predictions by botanical countries to obtain standardised predictions:

$$Prop_{k}=\frac{1}{n_{k}}\sum_{i=1}^{n_{k}} S_{i,k}\left( t \right) S_{i,k}\left( t \right)=exp\left( -\int_{t_{0}}^{t} h_{i,k}\left( t|X=x \right) dt \right), i=1,...,n_{k}$$

where, *Prop_k_* is the expected proportion of species still awaiting description or geolocation in botanical country k, *S_i,k_(t)* denotes the probability that species i awaits description or geolocation in k at time t, *n_k_* denotes the number of species used for model fitting in k.
