## Supplementary material for "Plant diversity darkspots for global collection priorities": Figure S1

**Fig. S1** Principal Component Analysis of 17 sustainable development goals (SDGs) and distribution of botanical countries along PC1 and PC2. All are coloured by income group as delineated by the United Nations: high income (HIC), low income (LIC), lower middle income (LMIC) and upper middle income (UMIC). The SDGs measure a country’s progress towards achieving the following: (1) no poverty, (2) zero hunger, (3) good health and well-being, (4) quality education, (5) gender equality, (6) clean water and sanitation, (7) affordable and clean energy, (8) decent work and economic growth, (9) industry, innovation and infrastructure, (10) reduced inequalities, (11) sustainable cities and communities, (12) responsible consumption and production, (13) climate action, (14) life below water (15) life on earth, (16) peace, justice and strong institutions and finally (17) partnerships for the goals

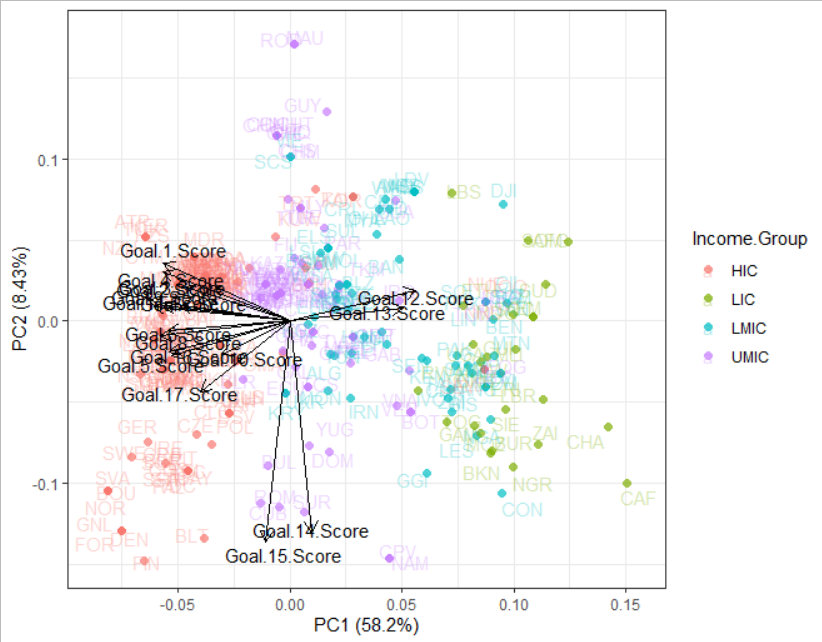
