## Supplementary material for "Plant diversity darkspots for global collection priorities": Figure S2

**Fig. S2** In panel **(a),** the Linnaean shortfall (blue gradient) is quantified as the predicted number of species yet to be described scientifically based on the estimated probability of species to remain undescribed by 2021. The Wallacean shortfall (red gradient) is quantified as the predicted number of species yet to be geolocated based on the estimated probability of species to have no geographic record available in a given botanical country by 2021. Colours indicate the relative magnitude of each shortfall, with light grey and dark brown colours representing small and large values for both shortfalls, respectively. In panel **(b),** the top 5 botanical countries with the largest combination of both Linnaean and Wallacean shortfalls are reported by continent. Predictions were rescaled between 0 and 1 for comparison.

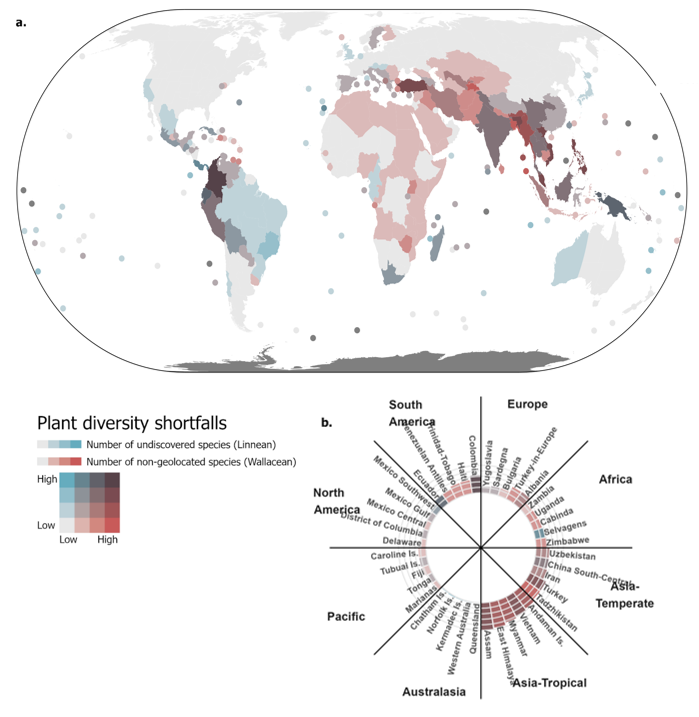
