## Supplementary material for "Plant diversity darkspots for global collection priorities": Figure S3

**Fig. S3** Panel **(a)** maps the year of peak description rate. Older peaks are coloured in brown and recent peaks in cyan. Taxonomic species descriptions have peaked in the 1800s in Europe, while they are peaking now in North Africa, the Middle East and South Asia. Panel **(b)** shows the description rate for the period 2010-2019 with the top five botanical countries: Kriti, East Aegean islands, Mozambique Channel islands, Comoros, and New Caledonia.  Similarly, panel **(c)** maps the year of peak geolocation and panel **(d)** highlights geolocation rates between 2010-2019 with top 5 botanical countries: Colombia, China South-Central, Ecuador, Turkey and Tanzania.

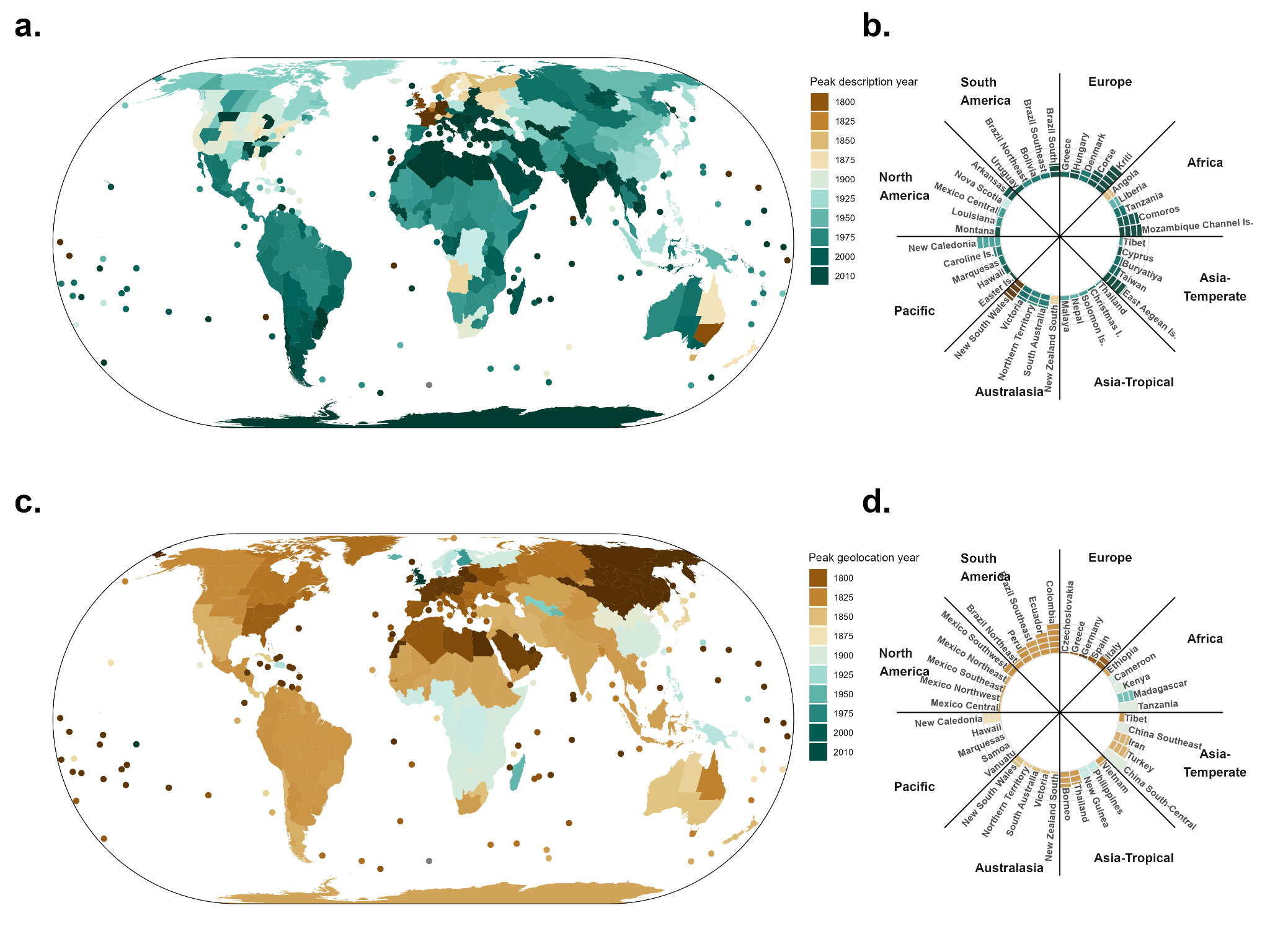
