## Supplementary material for "Plant diversity darkspots for global collection priorities": Figure S4

**Fig. S4** For both plots, dots represent botanical countries. In panel **(a)**, the x-axis represents the log-transformed number of species left to be described (scaled to a fixed area of 10,000 km^2^) and the y-axis represents the log-transformed average species description rate across 2010s (scaled to a fixed area of 10,000 km2). No relationship means there is little collection effort going on where there is more left to describe. In panel **(b)**, the x-axis represents the log-transformed number of species left to be geolocated (scaled to a fixed area of 10,000 km^2^) and the y-axis represents the log-transformed average species geolocation rate across 2010s (scaled to a fixed area of 10,000 km2). Positive relationship means sampling is currently happening in areas where there is more to geolocate.

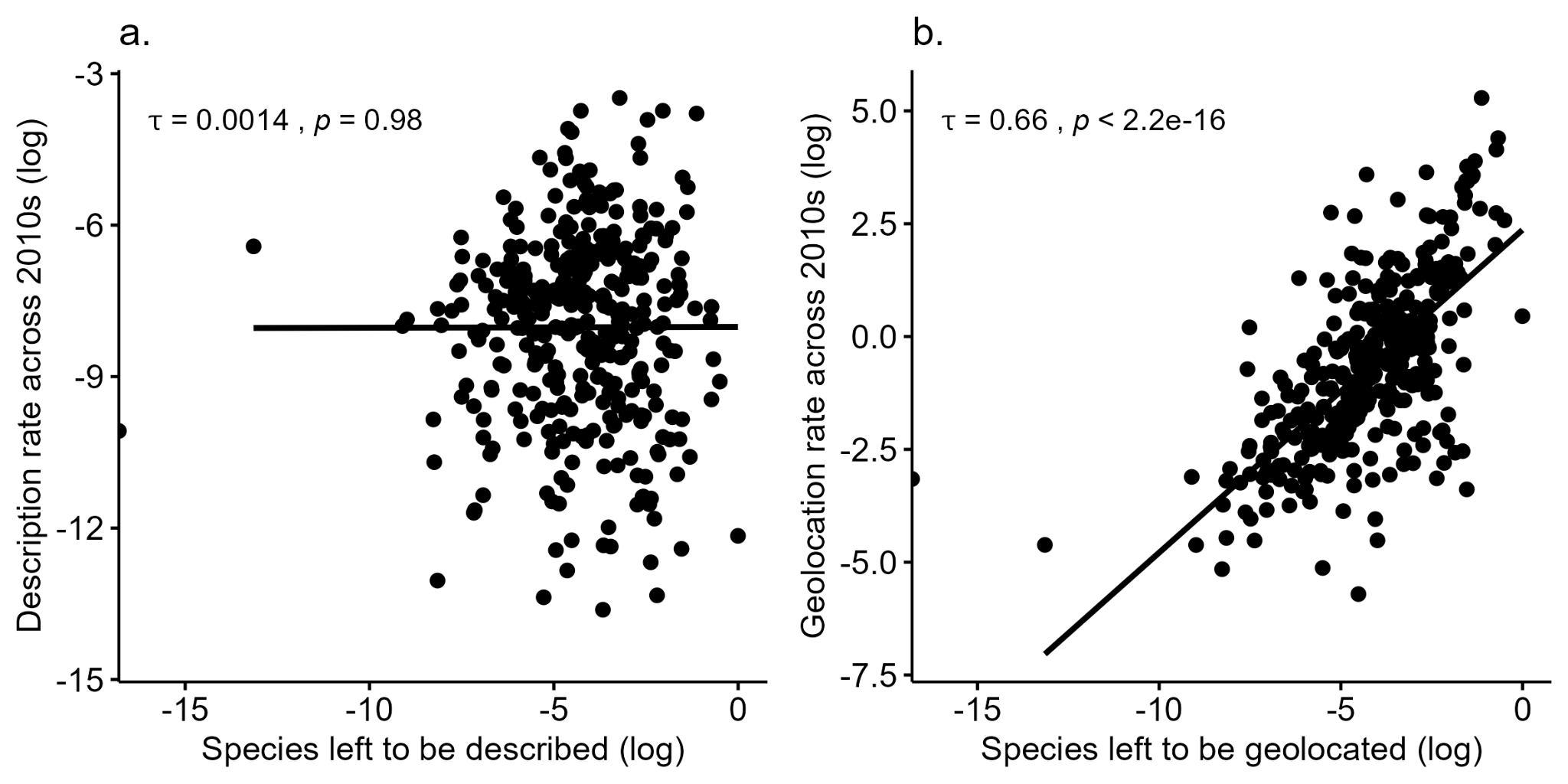
