## Supplementary material for "Plant diversity darkspots for global collection priorities": Figure S5

**Fig. S5** Dark red indicates botanical countries that are identified as darkspots and that contain hotspots. Black indicates botanical countries uniquely defined as darkspot (i.e. New Guinea). Orange indicates botanical countries containing hotspots but not overlapping with any darkspot (note that the area of hotspots is smaller than that of most botanical countries, and grey indicates areas containing neither darkspots nor hotspots.

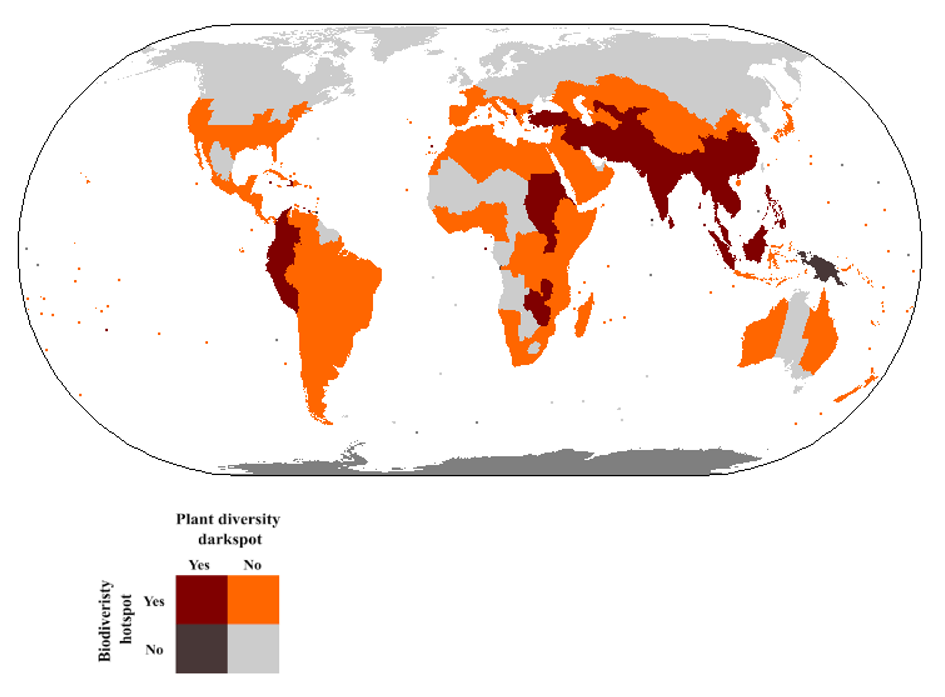
