## Supplementary material for "Plant diversity darkspots for global collection priorities": Figure S6

**Fig. S6** Collection priorities are displayed at both global and continental scale. Different scenarios investigate different tradeoffs between data gaps (baseline calculated from the combined linnean and wallacean darkspots scaled to a fixed area of 10,000 km^2^), income (PC1) and environmental protection (PC2) levels. In the top maps, we can see the top global (blue) and continental (green) collection priorities across scenarios, as well as their overlap (black) or absence (grey). The bottom matrix allows us to look at these scenarios in more detail, to see which countries are selected in which scenario.

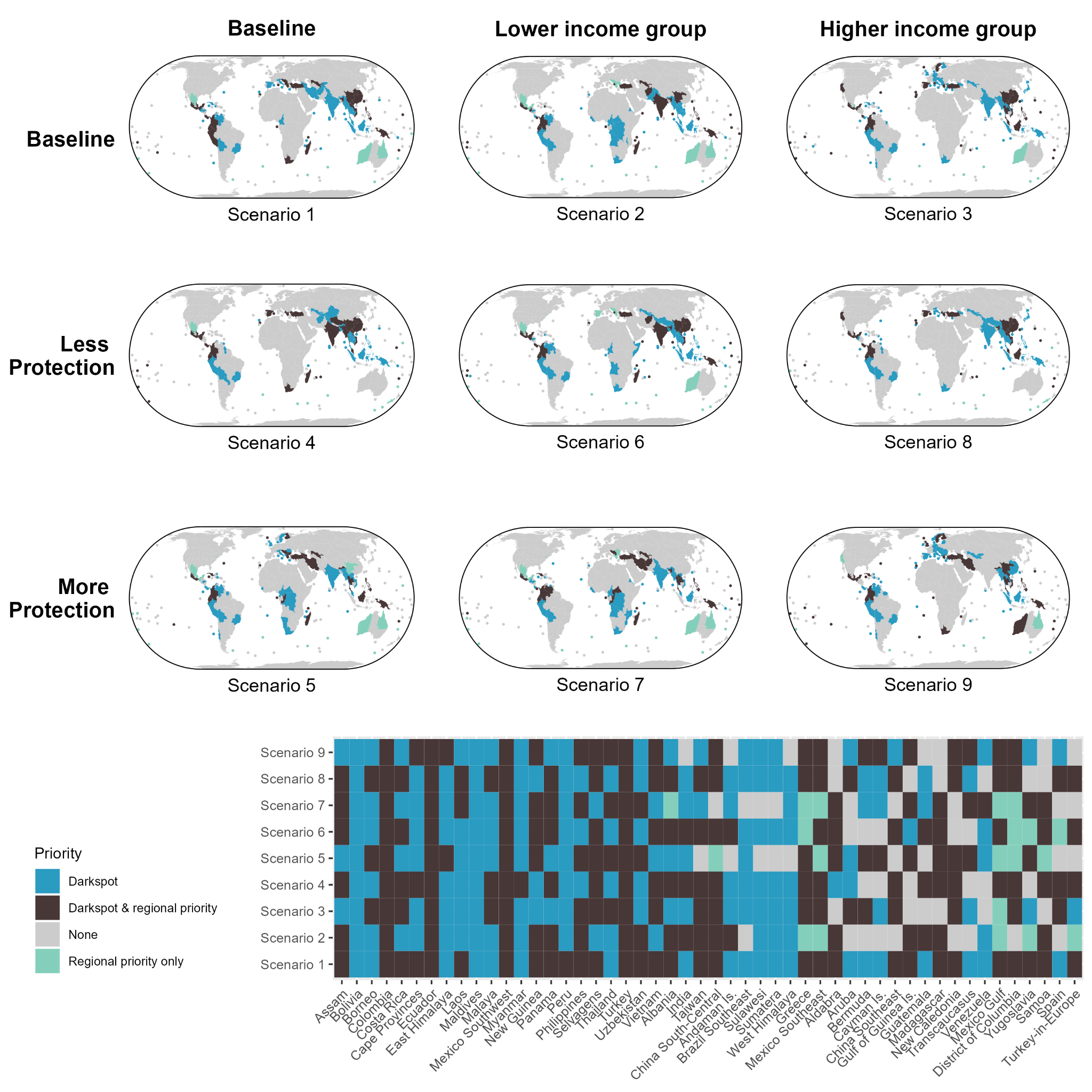
