## Supplementary material for "Plant diversity darkspots for global collection priorities": Figure S7

**Fig. S7** Botanical countries most likely to describe the most species taxonomically and geographically by 2021 and with the least capacity to collect these data (i.e., high PC1 Score). Panels **(a)** and **(b)** are primary estimates while c and d are scaled to a fixed area of 10,000 km^2.^ Panels **(a)** and **(c)** represent the geographic maps of the estimates while **(b)** and **(d)** represent the roseplot of the top 5 botanical countries with the largest shortfalls per continent.

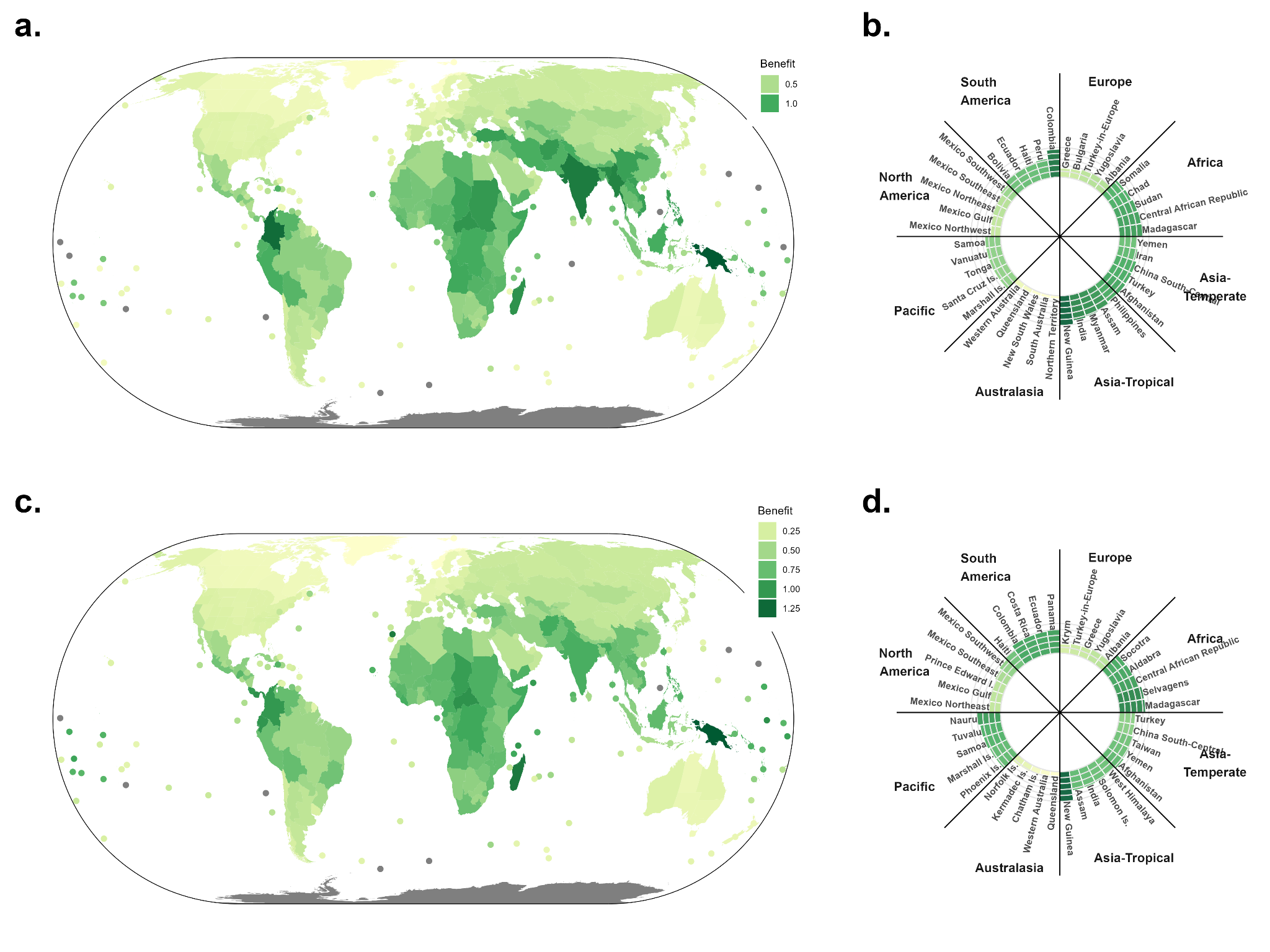
