## Supplementary material for "Plant diversity darkspots for global collection priorities": Figure S8

**Fig. S8** Botanical countries most likely to describe the most species by 2021 with the best capacity to collect these data (i.e., low PC1 score). Panels **(a)** and **(b)** are primary estimates while c and d are scaled to a fixed area of 10,000 km^2^ . Panels **(a)** and **(c)** represent the geographic maps of the estimates while **(b)** and **(d)** represent the roseplot of the top 5 botanical countries with the largest shortfalls per continent.

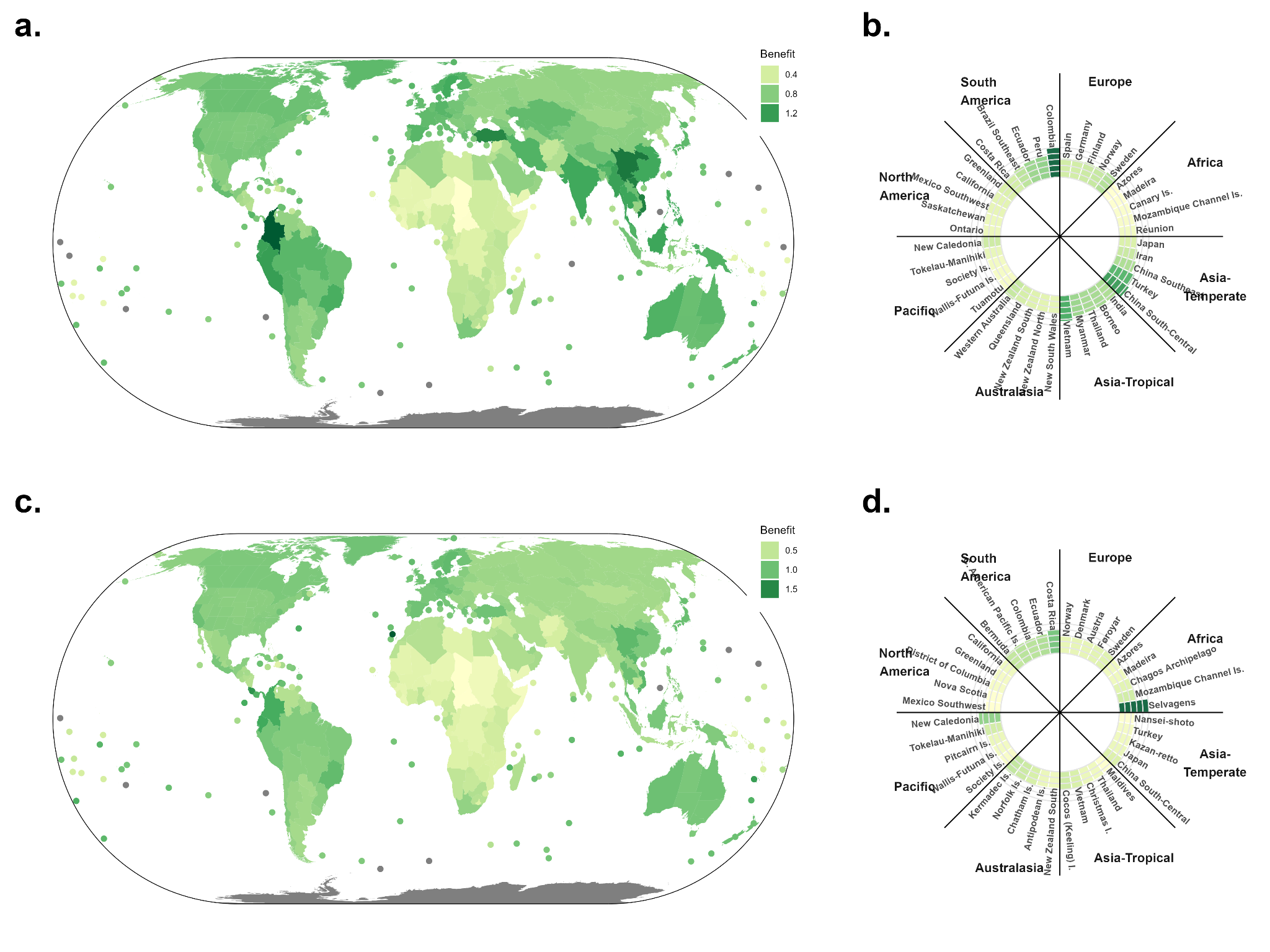
