## Supplementary material for "Plant diversity darkspots for global collection priorities": Figure S14

**Fig. S14** Botanical countries most likely to describe the most species by 2021 with the most capacity to collect these data and the most environmental protection (i.e., low PC1 and PC2 scores). Panels **(a)** and **(b)** are primary estimates while c and d are scaled to a fixed area of 10,000 km^2^. Panels **(a)** and **(c)** represent the geographic maps of the estimates while **(b)** and **(d)** represent the roseplot of the top 5 botanical countries with the strongest weight per continent.

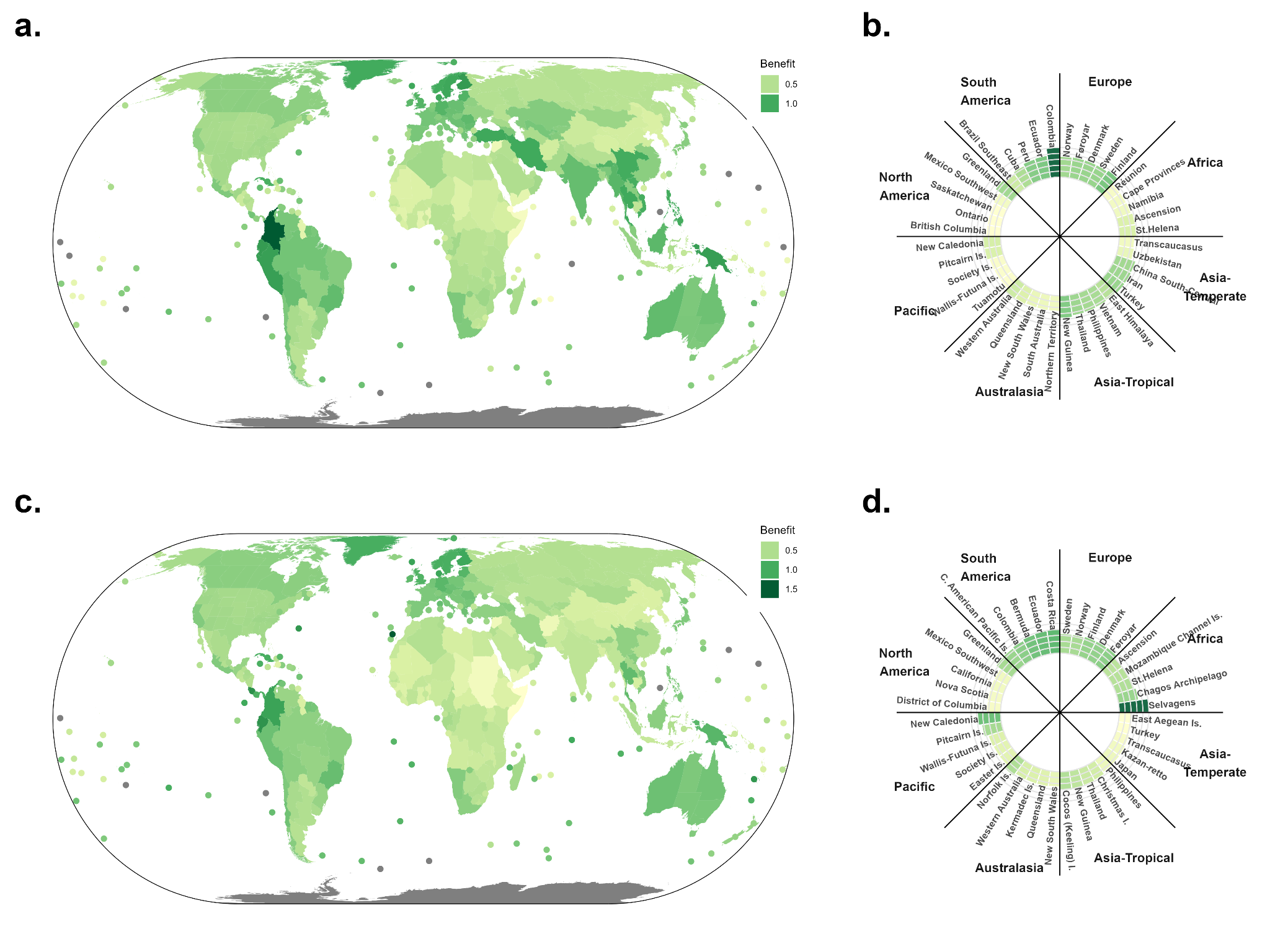
