## Supplementary material for "Plant diversity darkspots for global collection priorities": Figure S22

**Fig. S22.** Average expected benefit of countries when priorities for scenario 8 are recalculated 10,000 times using the mean, standard deviation, upper and lower limits of the darkspot index. Panels (a) and (b) are primary estimates while (c) and (d) are scaled to a fixed area of 10,000 km2. Panels (a) and (c) represent the geographic maps of the estimates while (b) and (d) represent the roseplot of the top 5 botanical countries with the largest shortfalls per continent.

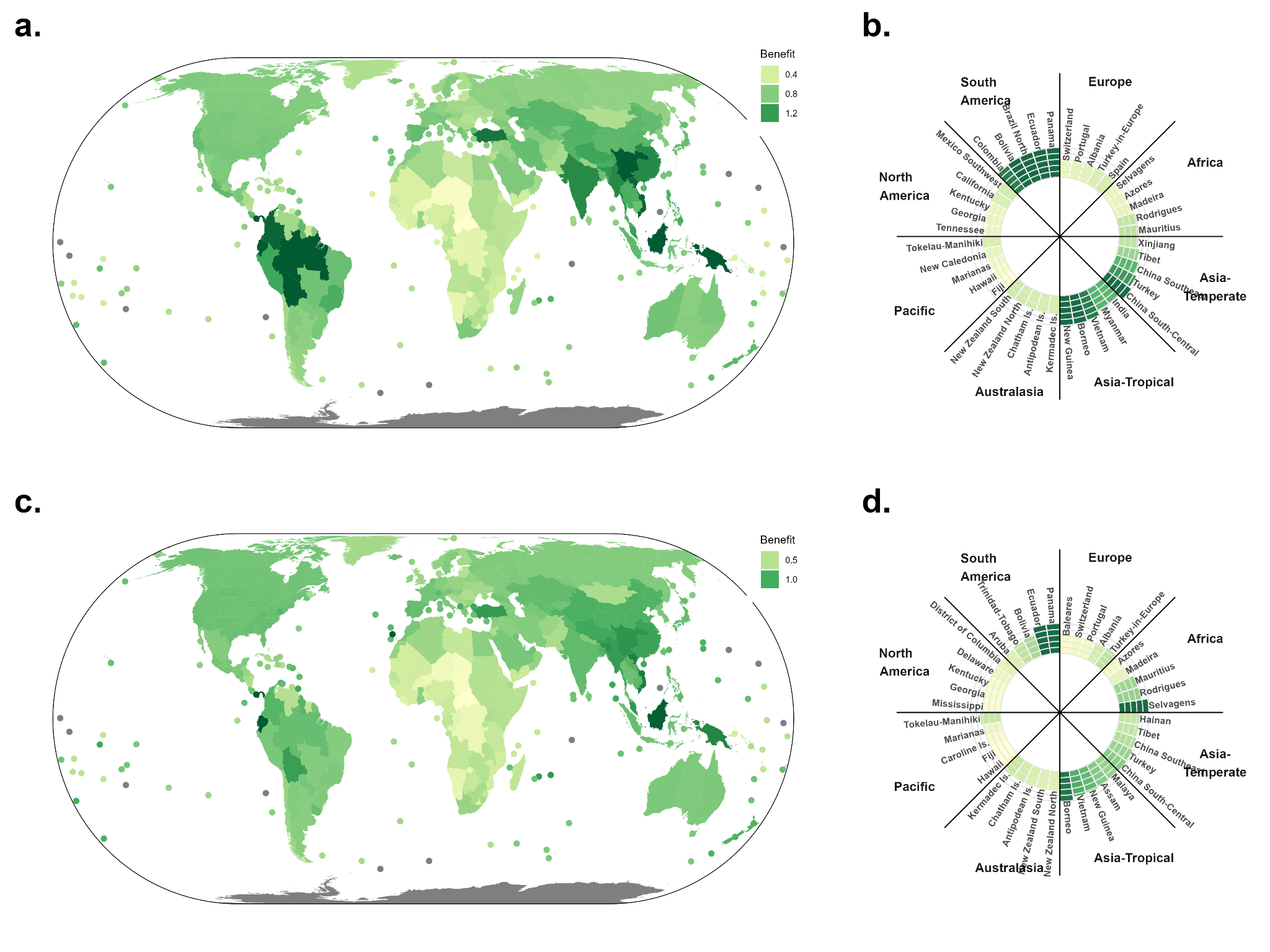
